# PTF-Vāc: An explainable and generative deep co-learning encoders-decoders system for *ab-initio* discovery of plant transcription factor binding sites

**DOI:** 10.1101/2024.01.28.577608

**Authors:** Sagar Gupta, Jyoti, Umesh Bhati, Veerbhan Kesarwani, Akanksha Sharma, Ravi Shankar

**Affiliations:** Studio of Computational Biology & Bioinformatics, The Himalayan Centre for High-throughput Computational Biology, (HiCHiCoB, A BIC supported by DBT, India), Biotechnology Division, CSIR-Institute of Himalayan Bioresource Technology (CSIR-IHBT), Palampur (HP), 176061, India; Academy of Scientific and Innovative Research (AcSIR), Ghaziabad, Uttar Pradesh-201002

**Keywords:** Transcription factor, Transcriptional regulation, Deep Learning, DenseNet, Transformers, Encoder-Decoder

## Abstract

Discovery of transcription factors (TFs) binding sites (TFBS) and their motifs in plants pose significant challenges due to high cross-species variability. The interaction between TFs and their binding sites is highly specific and context dependent. Most of the existing TFBS finding tools are not accurate enough to discover these binding sites in plants. They fail to capture the cross-species variability, interdependence between TF structure and its TFBS, and context specificity of binding. Since they are coupled to predefined TF specific model/matrix, they are highly vulnerable towards the volume and quality of data provided to build the motifs. All these software make a presumption that the user input would be specific to any particular TF which renders them of very limited use for practical applications like genomic annotations of newly sequenced species. Here, we report an explainable Deep Encoders-Decoders generative system, PTF-Vāc, founded on a universal model of deep co-learning on variability in binding sites and TF structure, PTFSpot, making it completely free from the bottlenecks mentioned above. It has successfully decoupled the process of TFBS discovery from the prior step of motif finding and requirement of TF specific motif models. Due to the universal model for TF:DNA interactions as its guide, it can discover the binding motifs in total independence from data volume, species and TF specific models. In a comprehensive benchmarking study across a huge volume of experimental data, it has outperformed most advanced motif finding deep learning (DL) algorithms. With this all, PTF-Vāc brings a completely new chapter in *ab-initio* TFBS discovery through generative AI.

**Short Summary:** The discovery of transcription factor binding sites (TFBS) in plants is challenging due to high variability across species and context-specific interactions. Traditional tools rely on predefined models and often fail in their cross-species applications. PTF-Vāc has implemented generative a universal deep-learning model that decouples TFBS discovery from predefined motifs, enabling accurate, species-independent TFBS/motif identification while outperforming existing methods by huge leads.

## Introduction

Transcription factors (TFs) play a crucial function in regulating gene expression at the initial stage of transcription by binding specifically to short DNA sequences across the promoter regions. The DNA binding sites, which usually consist of approximately 5 to 20 base pairs (bp) (Stewart et al., 2012; Mitchell et al., 1989), act as the recognition sites for the TFs, allowing them to control the transcription machinery (Davidson et al., 2003; Jones et al., 2020). Across the genome, these short, repeating and over-represented sequences define the binding motif for any given transcription factor. Thus, in order to decipher the control of gene expression, it is necessary to identify these binding spots or motifs. Recent technologies like chromatin immunoprecipitation (ChIP) (Ho et al., 2011), tiling arrays (ChIP-on-Chip) (Waldminghaus et al., 2010), and next-generation sequencing (ChIP-Seq) (Johnson et al., 2007) have enabled direct genome-wide identification of regions bound by any given TF. ChIP experiments allow for motif finding for any particular TF, as the regions obtained from ChIPs are much larger than the actual transcription factor binding sites (TFBS) while their TF specificity work as the guide to decode such binding motifs for any given TF (Zambelli et al., 2013). However, it is not feasible to perform ChIP experiments for all TFs, for every possible conditions and species, as the cost and possible search space shoots exponentially and almost infinitely. Thus, the computational models for TF:DNA interactions become an indispensable need.

At present, almost every TFBS discovery problem is preceded by motif discovery problem, and this has many issues. In current landscape of motif search, numerous challenges persist, reflecting the intricate nature of deciphering regulatory elements in genomic sequences. TF specific profiles and position weight matrices (PWM) have been age old and preferred approach where the TF binding regions (TFBRs) are excavated for possible alignments, based on which a scoring matrix is developed. The problem can be formalized as combinatorial optimization process which involves finding the combination of oligos that build the highest-scoring profile by exploring the search space of all possible combinations. Nearly all combinatorial optimization techniques, such as greedy, local search, stochastic search, and genetic algorithms, have been tried and applied over the years (Fan et al., 2008; Bucher et al., 1990). The introduction of PWMs, brought a quantitative dimension to motif representation, allowing for probabilistic assessments of TF binding (Stormo et al., 1982). Subsequent milestones included unsupervised statistical learning and clustering approaches, most popularly Gibbs sampling, Expectation maximization, and information theory based approaches (Lawrence et al., 1993). The creation of the databases like TRANSFAC and JASPAR offered the prime repositories of motifs from the experimental binding data (Matys et al., 2006; Sandelin et al., 2004).

A direct approach is an exhaustive search in which all possible motif candidates are scanned and guaranteed to find the most desirable one. However, such a direct approach frequently become unfeasible due to the exponential nature of the problem, resulting in highly demanding processing needs (Alipanahi et al., 2015; Trabelsi et al., 2019; Yang et al., 2019). On the other hand, statistical models, with tools like Consensus (Hertz et al., 1999), MEME (Bailey et al., 2009), MEME-ChIP (Ma et al., 2014), and AlignACE (Roth et al., 1998) as the prominent examples, make use of probabilistic approach for motif identification in DNA sequences. Among them, MEME-ChIP integrates three different algorithms to handle shorter and sparser motifs, allowing it to outperform other traditional TFBS discovery software tools (Ma et al., 2014; Jyoti et al., 2024). These algorithms have some drawbacks despite their effectiveness. Their sensitivity to user defined parameters is one such drawback, since poor decisions might affect the precision and effectiveness of motif recognition (Bailey et al., 2009). Furthermore, as demonstrated by the exploration-exploitation trade-off difficulties, these algorithms may have trouble with local optima, producing results that are not ideal (Hashim et al., 2019; Ma et al., 2014). An additional issue is that the solutions have limited biological interpretability (Kato et al., 2007).

Later on, with the rise of machine learning (ML) revolutions, motif discovery also witnessed its application with tools like SVMotif (Kon et al., 2007), which was developed to distinguish between TFBSs and non-binding sites by employing support vectors. Because of its adaptability, it was able to incorporate a variety of biological information and anticipate binding sites in sequences that were not yet been identified. However, the shortcomings of such ML approaches include their reliance on labeled data, sensitivity towards parameters, restricted biological interpretability, possible scaling problems on large genomic datasets, and their inability to understand more complex and hidden features. They rely exclusively on the input data features supplied by the user and do not attempt to extract more information or hidden relationships. As a result, the success of a ML model heavily depends upon the expertise and insight of its developer. Advanced ML approaches, notably exemplified by the arrival of DeepBind in 2015 (Alipanahi et al., 2015), signified a paradigm shift towards leveraging deep learning (DL) for motif finding. DeepBind looks for more comprehensive sequence contexts due to its emphasis on sequence information. It may need to be trained for use with data of different organisms due to its innate dependency upon species-specific training. In DL-based tools, the motif discovery stage is closely linked to the TFBR discovery stage. As a result, the accuracy of motif discovery is heavily influenced by the amount of noise and false identifications encountered during the earlier stages. Despite this, today DeepBind stands as the backbone philosophy for most of the TFBS discovery tools employing deep learning.

The variability in motif length and composition, coupled with context-dependent functionality, presents challenges in accurately identifying relevant motifs and binding sites. Addressing noisy data and distinguishing genuine motifs from false positives is essential (Ching et al., 2018). Furthermore, *ab-initio* motif discovery remains a complex problem, requiring algorithms capable of uncovering unknown patterns without prior knowledge (Bailey et al., 1998). Amid this all, there stands a big lacunae: a total ignorance towards stake of TF structure and its co-variability with the binding site. Very recently our group had conducted a comprehensive review and assessment of these software tools, addressing their shortcomings and proposing ways for improvement (Jyoti et al., 2024). The most common and gravest issue was their inability to address cross-species variation rendering them useful only for model species. As highlighted in Jyoti et al., 2024, plants exhibit significant genetic variability across their genomes, including in repeats, promoters, and TFs. Unlike animals, where GC content and repeat types are relatively stable, plant genomes show substantial variability. This variability affects TF binding sites (TFBS), making them more variable in plants than in animals which causes difficulties to address cross-species variations in plants and provide any credible results (Gupta et al., 2024; Jyoti et al., 2024; Kielbasa et al., 2005; Shiu et al., 2005; Bao et al., 2019; Lehti-Shiu et al., 2017; de Mendoza et al., 2013).

Understanding these issues, we had developed a novel approach, PTFSpot (Gupta et al., 2024), which learns from the interdependent variability of the TF’s structures and its binding regions to identify the most potential regions for any TF’s binding in total independence of any TF and species specific model (Gupta et al., 2024). The universal TF:DNA interaction model of PTFSpot makes it highly capable to determine the most potential binding regions for even unseen TFs and species. However, PTFSpot proposes a region of ∼150-162 bases and mimics DAP-seq like signals for most potential regions to look for the precise binding sites for any given TF. Unlike ChIP-seq, PTFSpot is not driven by limitations of experimental conditions. In turn, such proposed regions become highly reliable guide on which precise location of TFBS can be identified as well as can be used to build highly reliable motifs for any given TF, if the same structural and binding sites variation co-learning is carried forwards to deep-learning encoder-decoder system in a generative manner. The very same has been implemented in the present work through the developed software, PTF-Vāc. PTF-Vāc leverages from the highly refined information generated by PTFSpot to translate the longer binding regions proposed by PTFSpot into the most informative sites, the potential binding sequence/motif components using Transformer encoder-decoder for speech translation. Further to this, it applies interpretability by using Gradient Weighted Class Activation Mapping (Grad-CAM) (Selvaraju et al., 2017) to support the proposed motifs. Unlike any existing software, PTF-Vāc is a first of its kind auto-generative deep-learning system to decode precise binding sites of any transcription factor. The outcomes are groundbreaking, as PTF-Vāc consistently delivered the best performance, whether given extensive data or just a single sequence, putting it far ahead of the existing software pool for the same purpose. It has successfully decoupled the problem of motif finding and binding site discovery while working on the plant specific system.

## Results and Discussion

### Deep co-learning by Transformer-DenseNet system on the words and structure performs consistently well

Plant genomes are extremely complex, highly dynamic, with a greater diversity and complexity of repetitive sequences compared to animal genomes, which tend to be more stable in repeat content and composition (Shiu et al., 2005; Kielbasa et al., 2005; Bao et al., 2019; Lehti-Shiu et al., 2017; de Mendoza et al., 2013; Jyoti et al., 2024; Gupta et al., 2024). This variability in plant genomes means that the binding regions and motifs of a TF may differ significantly between species, even for the same TF, as has been highlighted by the above mentioned recent studies. These reports have brought attention to the prevalent under performance in the majority of existing TFBS discovery categories during TFBS annotations, often resulting in a high incidence of false positives (Jyoti et al., 2024). A potential solution lies in understanding the variations in TF structure and their corresponding binding sites. Recently, all of these concerns were raised and addressed by our group through the software PTFSpot (Gupta et al., 2024).

TFBRs discovery is closely tied to motif discovery, relying on precise TF-specific information from binding experiments like ChIP-seq. These methods assume that input sequences bind to a predefined TF, but they have several limitations: 1) Non-specific sequences can introduce noise, 2) It’s impractical to obtain TF-specific sequences experimentally due to high costs and vast search space across species, 3) Performance is highly dependent on the data volume, becoming weaker with fewer sequences and overly noisy with large datasets due to the inclusion of non-binding motifs. Moreover, these approaches are ineffective for genomic-scale scanning, especially in cases where no predefined guide exists, such as with novel genomes. In such scenario, the TF binding region proposed by PTFSpot becomes a highly accurate guide, on which suitable further learning could be performed and deep-learning decoders may be applied to speak out the precise TFBS motif candidates. In the PTF-Vāc implementation, as outlined in the methods section, Transformers and DenseNet system simultaneously acquired knowledge about TFBS, considering both sequence contexts and the protein structure of the TF. Taking this consideration into account, Dataset “E” from PTFSpot was retrieved, while focusing solely on positive instances as the problem in the present work was not classification but text generation (binding site generation). The associated 3D protein structure for each TF was also considered. The Transformers-DenseNet system underwent training and testing on a dataset comprising 48,000 instances of binding regions for the selected 40 TFs, employing a 70:30 split ratio with no overlap. The raised model attained an accuracy of 92.76% on the test dataset.

Now, a very interesting question arises that which features contributed the most in the observed performance. An ablation analysis was conducted, evaluating the performance of one after another considered properties, both individually and in combination. A total of six distinct sequence-derived word representations were utilized. The results indicated a clear trend, as more complex representations were introduced, significant increase in accuracy was observed: dimeric representations returned 32.56% accuracy, trimeric representation achieved 46.1%, tetrameric reached 59.51%, and pentameric representation attained 76.49% accuracy. This progression continued with hexameric and heptameric representations, which returned accuracy values of of 80.2% and 83.77%, respectively. It was evident from these findings that the contribution of higher-order words improved the performance mainly because of increase in the vocabulary size due to higher number of representational words. For instance, the pairing of pentameric and hexameric representations (totaling 311 words) resulted in a significant leap in the accuracy to 89.97%. The best instance of the additive benefits of combining representations was illustrated when incorporating pentameric, hexameric, and heptameric representations, resulting in an impressive accuracy of 92.76% with a total of 465 representative words **(Supplemental Table 1 Sheet 1)**. These results emphasize the necessity of information sharing among different representations, as they collectively harnessed their unique attributes to raise a better model.

Optimization of the hyper-parameter on the above mentioned model was implemented to refine the model further which yieled the reported accuracy of 93.2% on the test dataset. An exploration into the performance numbers of encoders and decoders for the Transformers-DenseNet system revealed that the sharpest improvement occurred at the sixth encoder-decoder layer, with performance stabilizing through the eighth layer before declining. As a result, the system was implemented with six encoder-decoder pairs. The final processing for text generation was handled by the decoder’s feed-forward layer. The final hyperparameters of the output layer of the optimized model were: {”Activation function”: softmax, “Loss function”: cross-entropy, “Optimizer”: Adam, “learning rate”: 0.0001, “Epochs”: 50, “Batch size”: 6}. **Figure 1** shows the detailed workflow and the implemented transformer architecture.

**Figure 1:**
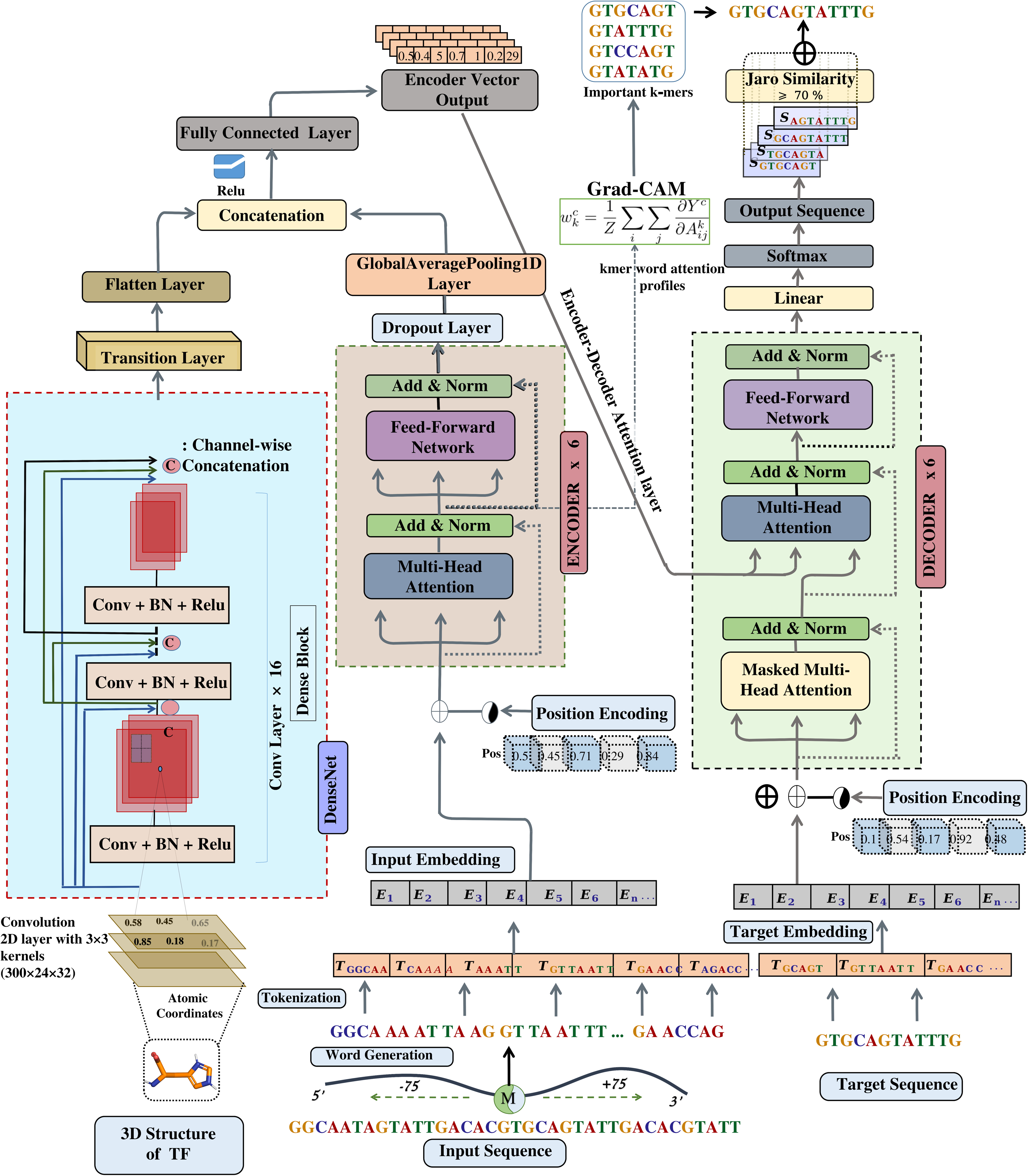
The PTF-Vāc Deep Co-learning system. Bi-modal deep-learning by Transformers-DenseNet Encoder:Decoder system was implemented to detect TFBS in TF binding regions derived from PTFSpot. The encoder processes binding region sequences represented in penta, hexa, and heptamer word representations, while the decoder receives the binding site information for the corresponding binding region. Six layers encoder:decoder pairs were incorporated. Simultaneously, a DenseNet consisting of 121 layers accepts input in the form of the associated TF’s structure. The learned representations from the encoder and DenseNet are merged and passed to the encoder-decoder multi-head attention layer, which also receives input from decoder layers. Subsequently, the resulting output undergoes conditional probability estimation, playing a pivotal role in the decoding process. Following layer normalization, the model proceeds to calculate the conditional probability distribution over the vocabulary for the next token. Finally, the resultant tokens were converted back into words, representing the binding site for the given sequence.

Afterwards, this was assessed that how much the observed performance gained from consideration of the TF’s structure while looking into the target binding region sequence. To assess that, the deep-learning system was raised again without the 3D structural information of the TF and the corresponding DenseNet parallel system which handled the structural information part using the same training dataset on which above models were trained. The resulting accuracy for the developed model dropped sharply to 73.9%, a significant drop. This provided strong support for the importance of simultaneously considering sequence and structural variability as previously demonstrated by PTFSpot too. AlphaFold2 generated structures were helpful to assess relative structural arrangements to create corresponding binding preference maps. An interesting observation was made when the 3D structural information was dropped from the model. Visible in the **Figure 2A**, two groups were formed “Highly Affected” and “Lesser Affected”. 19 TFs were “comparatively” lesser affected with an average of ∼12.6% accuracy loss whereas 21 TFs were affected more with an average accuracy loss of ∼21.23%. Results highlights that there were not a single TF which remain unaffected by the removal. Yet, since there was some sort of groupings based on the impact of structural information, we analyzed the secondary structure composition of these 21 affected and 19 less affected TFs to assess their structural differences. We studied the propensities for each amino acids within a TF towards Alpha-Helix, Beta-Strand, and Coil structures after normalization. A t-test was performed to compare the structural distributions between the two groups. The results showed a significant difference in the alpha-helix content (p-value = 0.0007), suggesting a potential structural differences. In contrast, Beta-Strand (p-value = 0.9797) and Coil (p-value = 0.15) showed no significant differences. The lesser affected group had higher share of alpha-helix, which makes sense. Alpha-helix helps in DNA motif recognition in major grooves (Garvie and Wolberger, 2001). Thus, a sequence motif information from DNA can compensate for the structural information loss to some extent. In terms of domains, there was no significant difference between the two groups.

**Figure 2:**
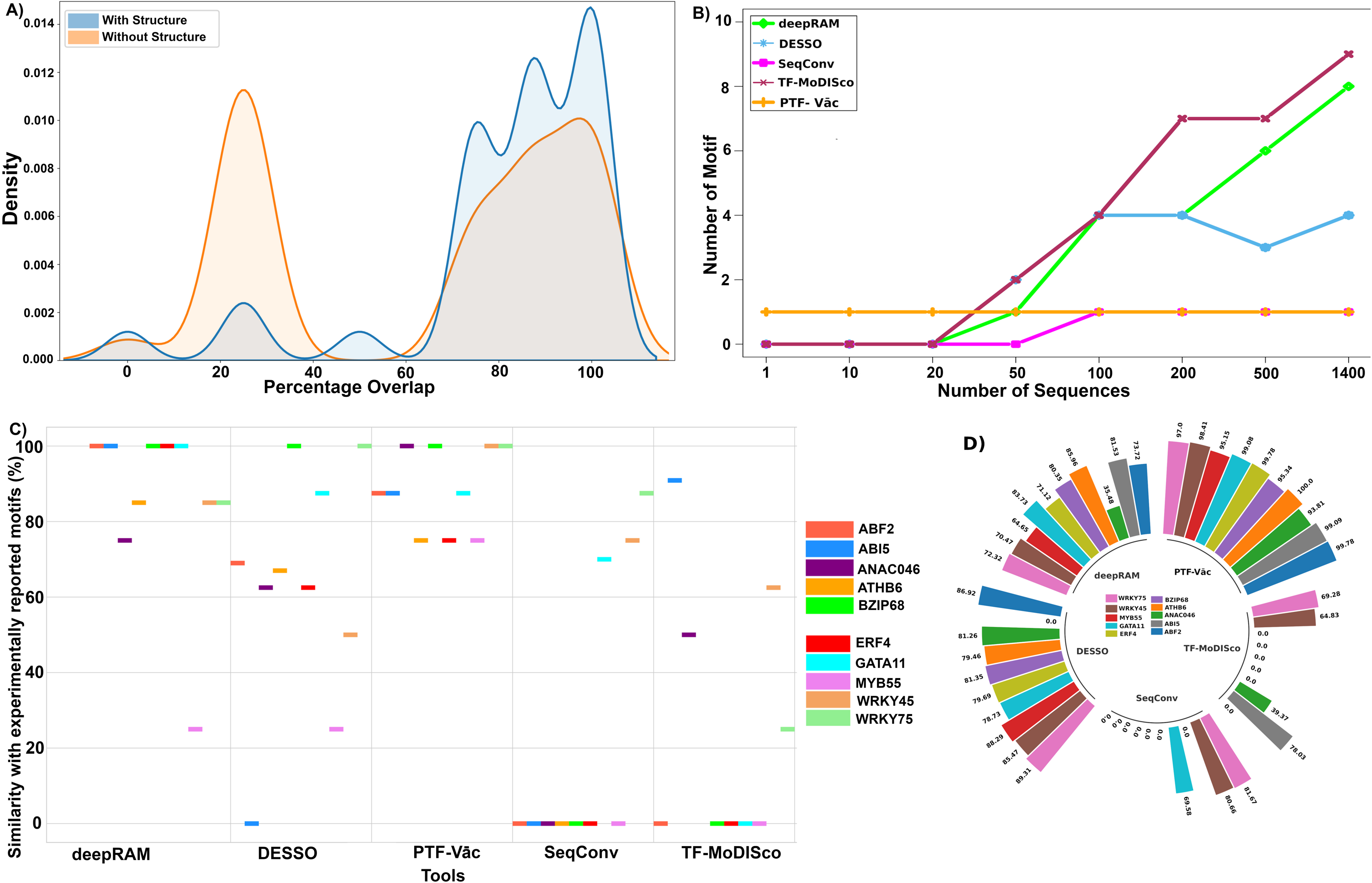
Performance benchmarking of the PTF-Vāc Deep Co-learning system. **A)** The performance of the PTF-Vāc over the test set with and without associated TF 3D structure. Incorporation of co-learning with TF structure played a major role in accurately finding the binding sites. Percentage overlap on the x-axis was calculated based on the overlap between the binding site identified by PTF-Vāc the actual binding site provided in the testing data for the same TF. This was done base-by-base using the following the equation (Number of common bases between the PTF-Vāc generated binding site and the binding site already provided in the testing data) / (Total number of bases of the binding site already provided in the testing data) × 100. **B)** Performance of different software on WRKY45 TF dataset of varying lengths ranging from one sequence to the whole dataset. Except PTF-Vāc, of the compared tools’ performance varied a lot with variation in the input volume. PTF-Vāc performance was totally unaffected by this, demonstrating its robustness and credibility. **C)** Comparative benchmarking showcasing the discovered binding sites’ similarity with the experimentally reported motifs for the selected 10 TFs for representation purpose. **D)** Comparative benchmarking showcasing the discovered motif data coverage for the 10 TFs (for representation purpose). For all the studied TFs, the details have been made available in the Supplemental Table 1-2.

### The identified binding sites displayed significant enrichment and matched the experimentally reported motifs

The next step was to determine the experimental binding data enrichment status of the reported binding sites by PTF-Vāc for any given TF as well as its closeness to the experimentally reported motifs. Experimental information on plant TFs, their binding sites, and interactions is far more limited than that of human TFs. Only for a few plant TFs experimentally validated motifs and matrices are available and for very limited number of species (Jin et al., 2017). Due to such constraints for experimentally validated motifs availability for plants, the present comparison was restricted to 30 TFs (DAP-Seq) for *A. thaliana* and six TFs (ARR1, FHY3, LEC2, LHY3, KAN1 for *A. thaliana* and GLYMA.08G357600 for *Glycine max*) with ChIP-seq data, having experimentally reported motifs at JASPAR (**Supplemental Table 1 Sheet 2**). The six TFs belonged to the families that were never been used during training of the model. These reported motifs ranged from 8 to 15 bases. The complexity of plant genomes, species diversity, and the resource-intensive nature of experimental validation are just a few of the obstacles that might make data accessibility in the context of plant genomics a limiting factor. This has to be noted that the experimentally retrieved motifs for these 36 TFs were not part of either the training or the testing datasets.

In order to ensure a uniform motif search approach for a fair comparison, these motifs underwent scanning using a methodology similar to that utilized in the PTFSpot approach (Gupta et al., 2024). Subsequently, the binding sites reported by PTF-Vāc were evaluated for statistically significant enrichment in the corresponding binding data for the given TF. It was discovered that for all the studied TFs their identified motifs were significantly enriched (binomical test p-value << 0.01) for their respective binding region data. PTF-Vāc generated motifs covered at least 83.26% binding data and went up to 100% coverage of the binding data (**Supplemental Table 1 Sheet 3**).

The resemblance between the experimentally reported motif and its corresponding matching motif from PTF-Vāc was assessed using TOMTOM, a part of MEME suite (Bailey et al., 2009). This comparison aimed to assess the degree of similarity and consistency between motifs identified through experimental techniques and those obtained through PTF-Vāc. All motifs retrieved from PTF-Vāc in this study for the 36 TFs matched with the experimentally reported motifs from JASPAR (TOMTOM p-value: <<0.01) (**Supplemental Table 1 Sheet 3**). 100% similarity to the experimentally reported motifs was observed for PTF-Vāc generated motifs obtained for ANAC046, ANAC055, ANAC083, ANAC092, ARR1, BIM2, bZIP68, ERF13, GATA15, WRKY18, WRKY30, WRKY45, and WRKY75. The consensus motif T[TC]GCCGAC (for RAP2-10) was the least abundant among the PTF-Vāc motifs, covering 83.26% of the bound data. Similarly, the consensus motif [AG][AG]AT[AG]CGCAA[TA], [TC]AAT[CA]ATT, [AC]GAATCT[ACGT], and [TC]CAACGC[TG] were the most abundant motifs, covering 100% of the bound data for ANAC083, ATHB6, ARR1, and WRKY30, respectively. **Supplemental Table 1 Sheet 4** provides a detailed comparison between motifs reported in this study and their experimentally reported counterparts. This part of the study further validated the methodology of PTF-Vāc and its identified TFBS motifs.

### Explainable deep learning using Grad-CAM gives insight into the properties which determine TF:DNA interaction

As discussed in methods section, for the implementation of Grad-CAM in PTF-Vāc, we chose the normalization layer after the multihead attention layer (MHA) as the layer of interest **(Figure 1)**. This layer contains distribution maps for various sequence k-mers. Following the weighting method proposed by the Grad-CAM (Selvaraju et al., 2017), we quantified the importance of these sequence k-mers and computed a weighted summation of all the distribution maps. This aggregated map identifies key k-mers crucial for binding activities. We extracted 5-mers, 6-mers, and 7-mers from positive sequences. Each unique k-mer was assessed to determine if it showed enrichment among the highest average Grad-CAM scores associated with each TF. Our results using the built models recapitulate multiple known trends. For example, the top scoring k-mers for any given TF often closely matched with the reported core motif of that TF itself.

The next step was taken to determine how accurate Grad-CAM captured the context relationship to the binding site for a TF. We selected those 30 TFs (DAP-seq) on which enrichment benchmarking was done in this study as discussed above. For all the 30 TFs, the top scoring k-mers for the considered TFs often closely matched with the motifs reported by PTF-Vāc encoder-decoder (**Supplemental Table 1 Sheet 5**). For WRKY18, highest agreement of 84.94% was found between the PTF-Vāc encoder-decoder generated motif and Grad-CAM highest scoring region. While the lowest agreement of 60.82% was noted for ANAC046. Therefore, disagreement between Grad-CAM supported highest scoring area and Encoder-Decoder proposed scoring motifs ranged between ∼15-50%.

Therefore, it was necessary to locate these disagreeing results and check whether they agreed at lower ranked regions proposed by the Encoder-Decoder. Since, the Encoder-Decoder generates single TFBS motif candidates, while Grad-CAM provides scoring for every position, we looked in the Grad-CAM scoring rank for the regions identified as TFBS motifs by Encoder-Decoder. It was found that the Encoder-Decoder proposed TFBS motif ranged among 92.67% among the top 10 ranks of Grad-CAM scores (**Supplemental Table 1 Sheet 6**). Thus, the Grad-CAM interpretability emerged as a strong confidence measure and scoring for the proposed TFBS motifs. Besides this, the Grad-CAM score also provides ways to look into other promising TFBS motif regions. Next, we checked the rest top importance scoring profiles which did not match with the motifs reported from PTF-Vāc, with experimentally reported motifs from JASPAR. It was found that a good overlap existed between the high importance scoring regions and experimentally validated motif for all the remaining cases (value ranging from 35.47-97.02%) (**Supplemental Table 1 Sheet 5**).

Like for the sequence features, the structurally important amino acids in TF can also be evaluated for their importance. **Figure 3** illustrates how structurally important amino acids and binding regions sequence properties were identified, with case example of MYB88 in two compared species. Here, Grad-CAM was further applied to extract features from the 3D structures of the MYB88 TF for *A. thaliana* and *Z. mays*. Comparative structural analysis of TFs across species may elucidate evolutionary pressure and functional adaptations. It helped to explain what structural variations corresponded to the binding regions variations, when one moved across the species where shift in binding preferences of TFs was noted. Feature extraction was performed using the final convolutional layer of the build Transformer-DenseNet model. Grad-CAM identified few key regions within MYB88 in both species. Notably, these regions exhibited sequence variability, corresponding to the disorders regions **(Figure 3 AI-BI)**. Comparative analysis of these regions between *A. thaliana* and *Z. mays* revealed significant structural differences, with one disordered region identified in *A. thaliana* and three in *Z. mays*. Additionally, for MYB88 in both species, the top motifs were extracted with Grad-CAM and were compared with the PTF-Vāc generated binding sites. It was found that the identified a top scoring region from Grad-CAM and motif generated from PTF-Vāc were found overlapping **(Figure 3 AII-BII)**. Thus, Grad-CAM offered a promising avenue for reasoning the binding preferences of TFs. By highlighting important sequence regions and structural features it provides are additional independent confidence to the Encoder-Decoder identified TFBS motifs as well as suggests additional important regions found critical for TF binding.

**Figure 3:**
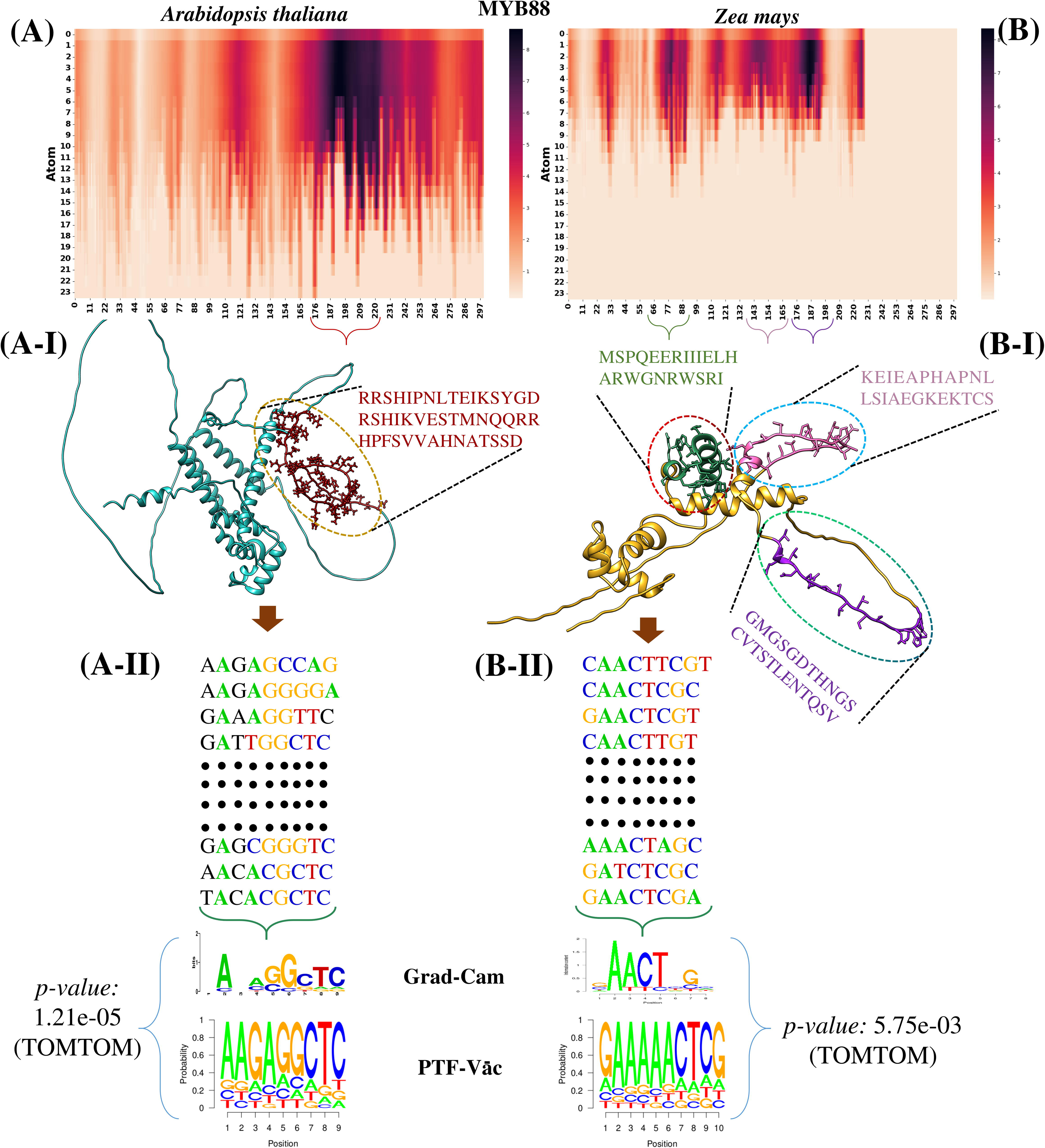
Explainability of PTF-Vāc Deep-Learning model. It helps to understand the responsible sequence and structural features for its output. Case of MYB88 comparison between *A. thaliana* and maize is discussed here. **A-B)** Grad-CAM based heatmap of MYB88 3D structure for *A. thaliana* and *Z. mays* respectively. **AI-BI)** Grad-CAM based heatmap revealed significant structural differences, with one disordered region identified in *A. thaliana* and three in *Z. mays*. **AII-BII)** Top sequence regions were was extracted for both species with Grad-CAM for TF MYB88. The explainability and importance scoring by Grad-CAM independently supported the identified binding sites by Encoder:Decoder. The highest scoring Grad-CAM regions were similar to the one proposed by Encoder:Decoder of PTF-Vāc.

### Comparative benchmarking and performance comparison

In order to get the final picture that how much performance leap PTF-Vāc approach offered over the existing state of the art methods for the same purpose, it was necessary to compare it with the most advanced software tools for objective benchmarking. All of the existing tools identify motifs from the sequences obtained from the binding experiments for some TF or require some guide where assumption is made that the user provides all sequences obtained from the binding data for some specific TF. If any diversion from this assumption is made, all these tools fail to generate quality motifs due to noise in their input data. As already mentioned above, all the existing methods have coupled motif discovery as prelude to binding sites discovery, and for all of them the above mentioned issue prevails. The same factor also leads to their limitations for cross-species use in plants where binding sites vary a lot along with TF structure (Jyoti et al., 2024; Gupta et al., 2024). This is also the cause that hardly any of the existing methods can credibly work for major jobs like genomic annotations for TFBS for any newly sequenced genome as none of them can work without a predefined motif matrix or PWM which they use for scanning. Expecting such motifs for novel genome wide scan itself is a point of mistake considering the variability across the plant species, as ably pointed out recently (Jyoti et al., 2024; Gupta et al., 2024). A possible solution is *ab-initio* binding site discovery which PTF-Vāc has achieved while basing upon the universal binding model of PTFSpot. It has learned the relationship between the variability of binding sites, TF structure, and context, making it free from any dependence on predefined motifs and matrices for any possible TF and any possible species. Thus, PTF-Vāc unlike the existing tools is capable to speak out the binding sites *ab-initio* even for single input sequence with the same level as it can do for multiple sequences. PTF-Vāc was compared to four recently published DL-based tools *viz.* deepRAM (Trabelsi et al., 2019), DESSO (Yang et al., 2019), TF-MoDISco (Shrikumar et al., 2018), and SeqConv (Shen et al., 2021) for different performance measure parameters which included: 1) Influence of volume of input sequence data on the binding sites discovery, 2) Overlap and similarity with experimentally reported motifs, and 3) Coverage of overall binding data given for any TF.

#### deepRAM and SeqConv

The approach employed by deepRAM deploy convolutional filters for feature extraction in motif identification with a focus on local pattern recognition. On the other side, SeqConv learns motifs from a deep CNN model by extracting active sequences in the rectification layer, yielding a 24-bp subsequence with the maximum convolution value for each sequence. However, their emphasis on individual filters may lead to redundant motif sequences and a limited capture of distributed motif representations.

#### DESSO

It employs convolutional filters to identify low-level features, followed by max pooling and fully connected layers for high-level feature extraction. Motif detectors learn patterns from the DNA sequences, and background sequences, which are selected to eliminate biases.

#### TF-MoDISco

It leverages contribution scores for assessing positional importance, offering insights into the significance of individual bases. It identifies high-importance windows called “*seqlets*”, clustering recurring similar “*seqlets*” into motifs after post-processing steps. While per-base importance scores aid in incorporating information from all network neurons, potential limitations arise when the model assign disproportionately high importance to specific positions based on prevalent motifs in the training data. This can lead to biases and potentially result in the overlooking of less frequent but biologically relevant motifs.

These four tools discussed above generate a PWM to discover motifs based on the learned patterns in the input data. However, in instances where these tools encounter noisy data during the learning process, they tend to produce ineffective motifs. The reliance on PWMs for motif prediction can lead to challenges in accurately discerning meaningful motifs. This emphasizes the importance of robust algorithms and noise-resistant strategies in the motif prediction process. In order to offer an unbiased performance comparison for each method, we cross-referenced the identified motifs with experimentally validated motifs obtained from the JASPAR database. We utilized available DAP-seq dataset for the 30 DAP-seq TFs for *A. thaliana* and six ChIP-seq TFs to scan and report their TFBS motifs across these sequences (**Supplemental Table 1 Sheet 2)**. As mentioned above, experimentally retrieved motifs for 36 TFs were not part of either the training or testing datasets used for model building and evaluation done previously.

### Influence of input dataset sizes on TF binding site discovery

In the first phase of our benchmarking, to measure the influence of input sequence data on the binding sites discovery, we conducted an in-depth assessment of the five software (deepRAM, DESSO, SeqConv, TF-MoDISco, and PTF-Vāc). Different input datasets based on number of sequences were constructed from the DAP and ChIP-seq dataset for every TF. The input this way ranged from one sequence to the whole dataset. It was observed for majority of TFs, deepRAM needed at last 50 sequences for motif identification, whereas DESSO and SeqConv required at least 100 sequences for motif discovery. On the other note, TF-MoDISco needed more than 200 sequences. Only PTF-Vāc discovered the binding sites irrespective of number of input sequences. Moreover, it was observed that deepRAM, DESSO, and TF-MoDISco often produced multiple motifs for a given TF as the volume of input sequences was increased, whereas SeqConv and PTF-Vāc consistently proposed a single TFBS motif, irrespective of input volume. **Figure 2B** provides a snapshot of the performance for the WRKY45 TF dataset for varying sizes, clearly showcasing PTF-Vāc’s ability to discover motifs regardless of input data size, in contrast to other software influenced by varying sequence data sizes. Detailed results of this analysis are presented in **Supplemental Table 1 Sheet 7**.

### Exploring similarity and overlap in TF binding site discovery

In the second phase of our benchmarking, we rigorously assessed the overlap and similarity between the motifs discovered by various software and experimentally reported motifs for all the 30 TFs (DAP-seq data) and six TFs (ChIP-seq data) as mentioned above. Using TOMTOM (Bailey et al., 2009), we evaluated the resemblance between experimentally reported motifs and their corresponding motifs discovered by each software. Notably, all the motifs identified by PTF-Vāc for the 36 TFs exhibited a significant match with experimentally reported motifs (TOMTOM p-value: <<0.01) **(Supplemental Table 1 Sheet 3).** Impressively, 100% similarity was observed between PTF-Vāc motifs and experimentally reported motifs for 13 TFs. Moreover, a minimum overlap of 83% was consistently observed for all TFs between motifs discovered through the PTF-Vāc and experimentally reported motifs. **Figure 2C** provides a snapshot of motif similarity between discovered motif and experimentally reported for different software tools for 10 TFs out of 36 for representation purpose only. For deepRAM, out of 36 TFs, only 24 TFs matched (TOMTOM p-value: <<0.01) with the experimentally reported motifs, whereas for DESSO, only 14 TFs matched with the experimentally reported motifs. deepRAM displayed a maximum overlap of 100% for the five TFs (ABF2, ABI5, bZIP68, ERF4, and GATA11) and minimum overlap of 25% for some of the TFs (CDF3, KAN1, and MYB55). Subsequently, DESSO displayed a maximum overlap of 100% for the three TFs (ATHB7, bZIP68, and WRKY75) and minimum overlap of 25% for one TF (MYB55). Only eight TFs for TF-MoDISco and four TFs for SeqConv matched with the experimentally reported motifs. Both, TF-MoDISco and SeqConv displayed the maximum data coverage of 90% with original dataset (**Supplemental Table 1 Sheet 4; Supplemental Table 2 Sheet 1-4**). In general, it was observed that most tools reported different motifs for the specified TFs, which did not closely match the experimentally reported motifs. Additionally, many tools generated multiple motifs due to the large input size. This all clearly exhibited much higher credibility of PTF-Vāc *ab-initio* approach than the compared tools and their approaches.

### Evaluating binding data coverage for each TF

In the third phase of our benchmarking, we conducted an exhaustive assessment of the overall experimental binding data coverage by the identified TFBS motifs for each TF. The results revealed that PTF-Vāc exhibited the highest data coverage for majority of the TFs with at least 83.26% data coverage and went up to the 100% coverage of the binding data for some of the TFs (ANAC083, ARR1, ATHB6, and WRKY30) (**Supplemental Table 1 Sheet 4**). Additionally, PTF-Vāc identified motif which achieved data coverage of 99% for the TFs ABF2, ABI5, ANAC058, ATHB15, ATHB7, BIM2, CDF3, ERF11, ERF4, ERF8, GATA11, GATA15, WRKY18, and WRKY25 (**Supplemental Table 1 Sheet 4; Supplemental Table 2 Sheet 1-4**). **Figure 2D** provides an overview of data coverage for discovered motifs for each software for 10 TFs. DESSO successfully discovered motifs for 34 out of 36 TFs, excluding ABI5, with a minimum data coverage of 23.46%. On the other hand, deepRAM discovered motifs for all 36 TFs, but 14 TFs exhibited a minimum data coverage of 75%. PTF-Vāc and deepRAM were the only tools successful in discovering TFBS motifs for all the 36 TFs. In contrast, TF-MoDISco failed to report motifs for several TFs (20 TFs). The motifs discovered for the remaining 16 TFs covered less than 90% of the binding data **(Supplemental Table 1 Sheet 4; Supplemental Table 2 Sheet 1-4)**. Another tool, SeqConv was capable to discover the motifs only for ARR1, GATA11, GLYMA.08G357600, KAN1, WRKY45, and WRKY75 with the data coverage of 37.48, 69.58%, 68.52, 52.53, 80.66%, and 81.68%, respectively. DL approaches for motif finding would benefit significantly from incorporating biologically relevant information, rather than focusing primarily on statistical and enrichment analyses (Jyoti et al., 2024). PTF-Vāc stands out as the leading solution for discovering relevant TFBS motifs in plants, far surpassing the second-best option, DESSO. The strong performance of DESSO can be attributed to its integration of biologically significant DNA shape features through deep learning, which many probabilistic models in other tools overlook. Thus, even for binding data coverage, PTF-Vāc emerged as the best one.

### Motif validation through molecular docking

We delved into examining the binding affinity between software (DESSO, SeqConv, deepRAM, TF-MoDISco, and PTF-Vāc) generated motif and the experimentally reported motifs associated with the same TFs using molecular docking. Randomly, five TFs were selected: ABI5, GATA11, MYB55, WRKY45, and WRKY75. Due to the general lack of protein 3D structure for these plant TFs, AlphaFold2 (Read et al., 2023) was used to model their 3D structure. Furthermore, the 3D structures of the identified binding sites were generated using the DNA sequence to structure web-server (http://www.scfbio-iitd.res.in/software/drugdesign/bdna.jsp). Molecular docking was executed employing pyDockDNA (Rodríguez-Lumbreras et al., 2022), and the obtained results were analyzed through Chimera (Pettersen et al., 2004) and PyMOL (Seeliger et al., 2010). Molecular docking has been used for such validation purpose in several studies investigating the interaction between TFs and DNA (Si et al., 2015; Konda et al., 2018; Mukherjee et al., 2015; Pandey et al., 2016; Zhu et al., 2014). The predicted model structures of the five TFs were docked with the binding sites identified by all the softwares and as well as experimentally identified binding sites for the same TF. The docking outcomes revealed that the binding affinities between the identified PTF-Vāc binding sites and their corresponding experimentally reported motifs for each TF were nearly similar. For instance, the binding affinities of GATA11 TF exhibited similar values for PTF-Vāc identified binding sites (docking score (ΔG): −111.33 kcal/mol) and the corresponding experimentally reported motif (docking score (ΔG): −111.1 kcal/mol). However, in the case of ABI5, the binding site derived from PTF-Vāc exhibited a higher affinity (docking score (ΔG): −76.6 kcal/mol) compared to the experimentally reported motif (docking score (ΔG): −67.43 kcal/mol). For other software (DESSO, SeqConv, deepRAM, and TF-MoDISco), the binding affinities of TF ABI5, GATA11, WRKY45, and WRKY75 exhibited lower values for identified binding sites by them and the corresponding experimentally reported motif. However, for MYB55, the binding site derived from DESSO exhibited a higher affinity (docking score (ΔG): −87.15 kcal/mol) compared to the experimentally reported motif (docking score (ΔG): −69.94 kcal/mol). But the identified binding sites by DESSO was not similar to the experimental one. The details of docking results are given in **Table 1**. Thus, the molecular docking study also supported the identified TFBS motifs from PTF-Vāc.

**Table 1:**
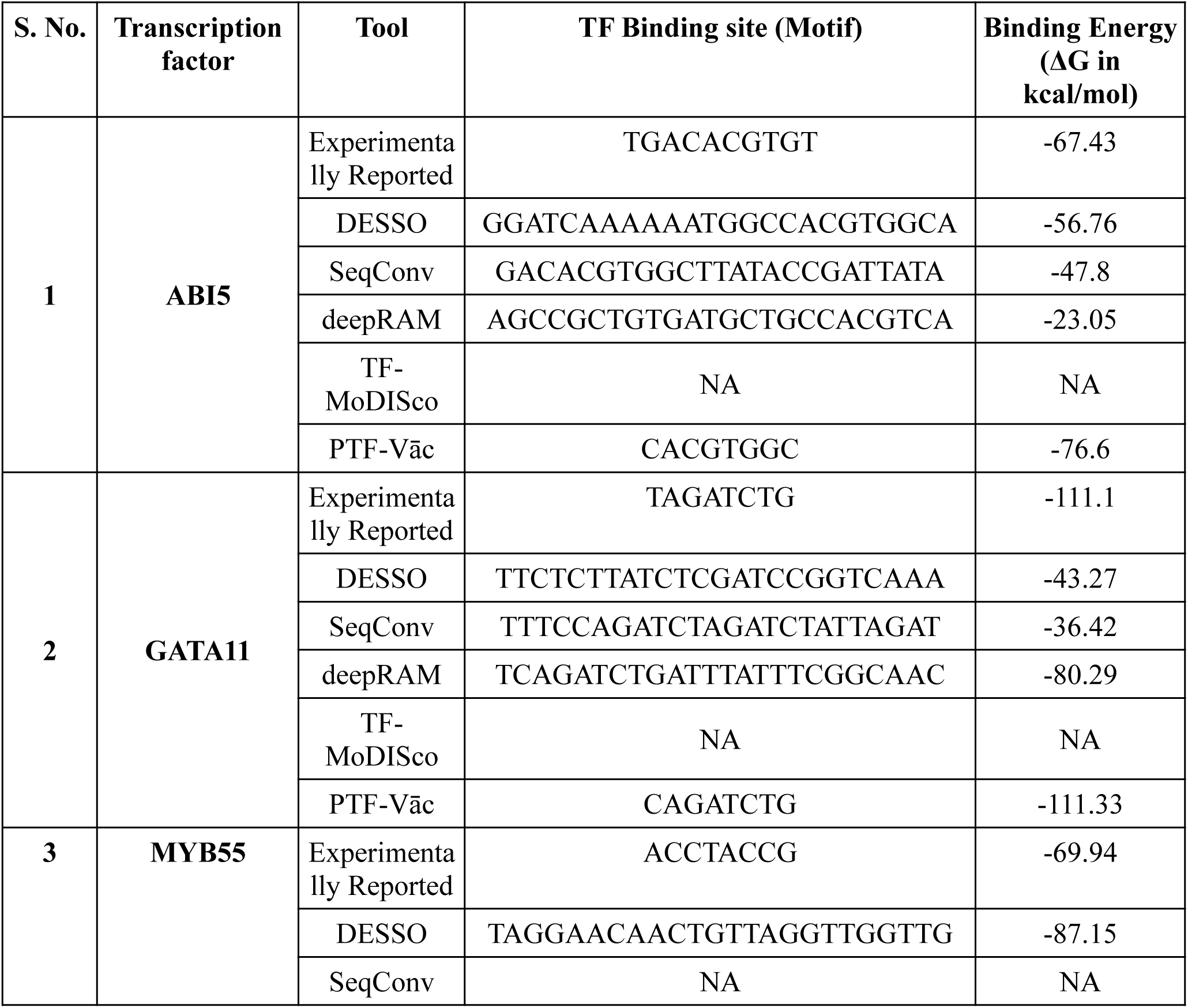
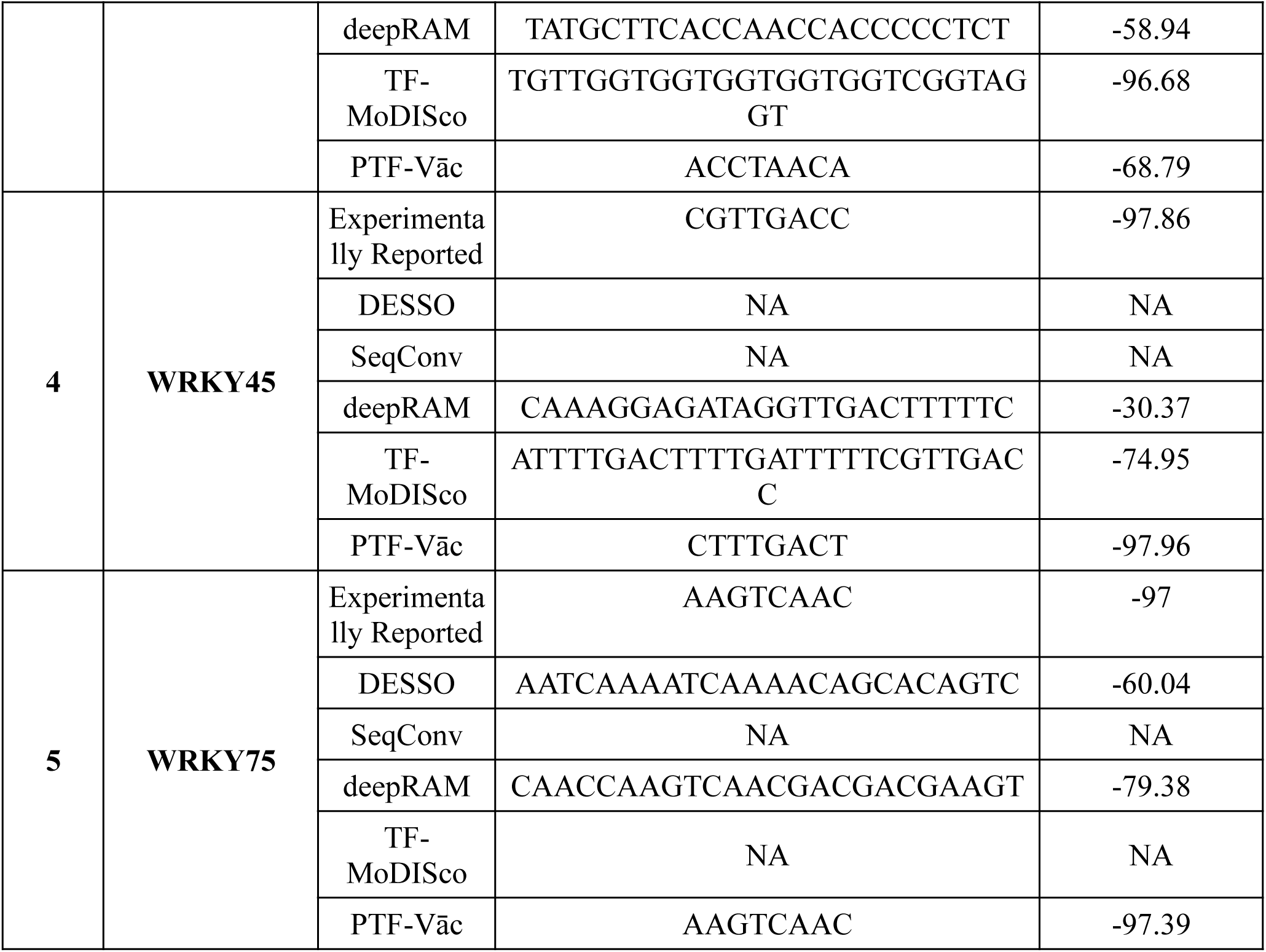
The details of docking results for five TFs for their bind*ing* motif validation through molecular docking for all the software. PTF-Vāc gave almost similar binding stabilizing energy as those observed for the experimentally reported binding sites. All of the reported binding sites formed highly stable complex with the corresponding transcription factor.

### PTF-Vāc accurately makes cross-species TFBS identification

The raised model’s final assessment was set to determine how well it performed to identify cross-species TFBS. In our investigation, we focused on *Zea mays* (seven TFs) and *Glycine max* (three TFs), for which ChIP-seq data and experimentally validated motifs were available. This set of TFs were never utilized before in this study and it was created solely for cross-species validation purpose for the model’s performance. The binding sites and motifs identified by PTF-Vāc demonstrated highly significant match (TOMTOM p-value << 0.01) with the experimentally reported motifs for all the considered TFs, affirming its efficacy in handling cross-species conditions. For bHLH47, EREB172, HB34, GLYMA-06G314400, and GLYMA-08G357600 highest match of 100% was observed with reported motifs. For three TFs, namely GLK1, MYBR17, and GLYMA-13G317000, the similarity of 88.9% was observed, whereas 77.8% of similarity was observed for EREB97 and TCP23. **Figure 4A** provides the comparison details for the identified binding sites by PTF-Vāc and corresponding experimentally validated binding sites for cross-species performance. The identified motifs covered maximum data above 90% for bHLH47, EREB97, EREB172, GLK1, HB34, TCP23, GLYMA-06G314400, and GLYMA-13G317000 (Supplemental Table 2 Sheet 5).

**Figure 4:**
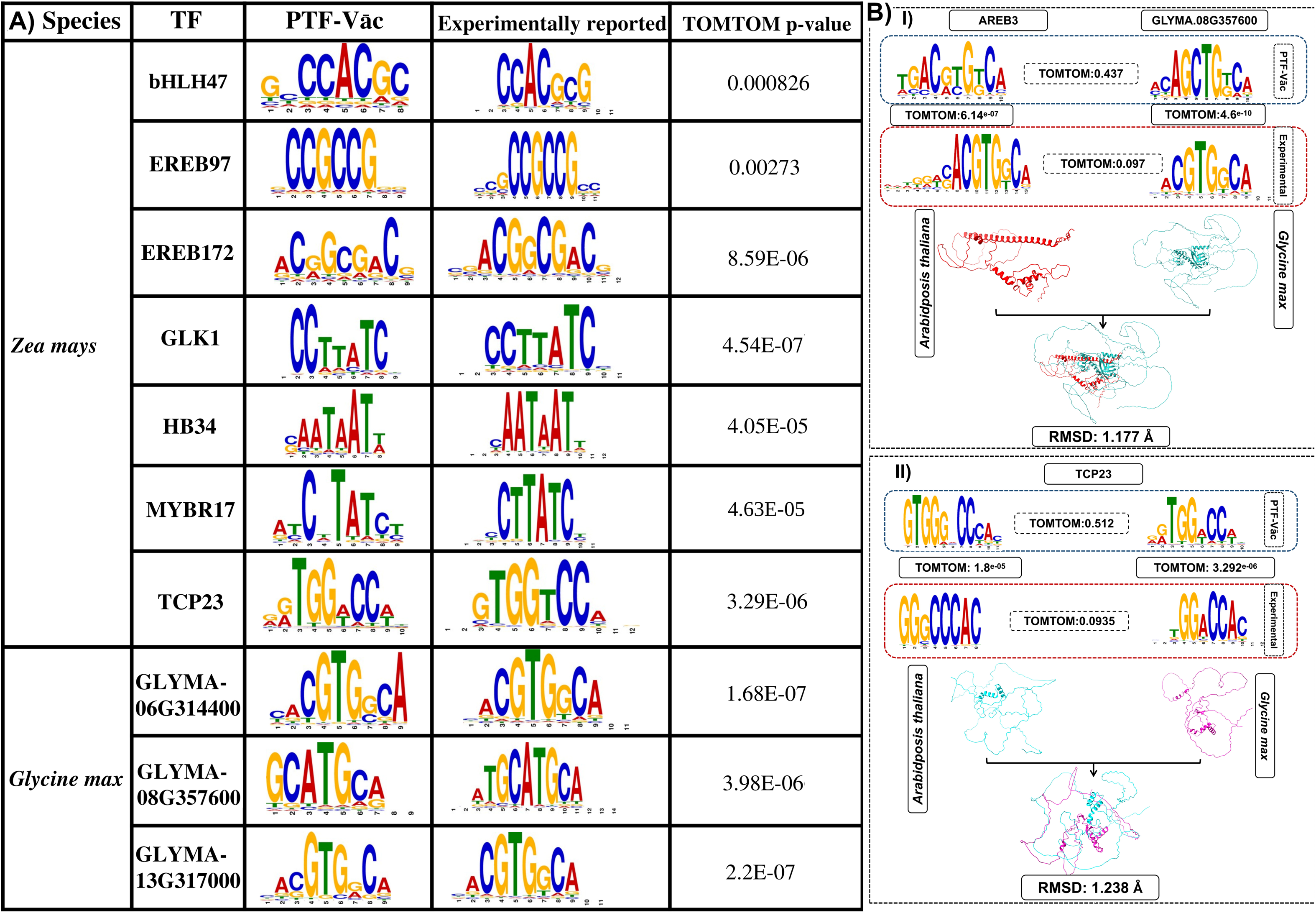
Cross-species performance of PTF-Vāc establishes it as the most reliable tool to identify TFBS across various plant species. **A)** Experimentally validated binding data and motifs from Zea *mays* and *Glycine max* were considered for test PTF-Vāc’s validity for its ability for cross-species performance. For all these 10 compared TFs and associated binding data, the binding sites and motifs reported by PTF-Vāc were found highly accurate and matching the experimentally reported sites with high significance when tested by TOMTOM. **B)** Comparison between experimental and PTF-Vāc binding motifs of TFs and their superimposed TF structures across species, with the structural differences measured in RMSD value, **(BI)** GLYMA.06G314400 between *A. thaliana* and *G. max*, **(BII)** TCP23 between *A. thaliana* and *Z. mays*.

Additionally, one more study was carried out to reconfirm that the algorithm actually learned the shift in binding patterns with structure and was not merely memorizing. The 3D TF structures as well as binding motifs of MYBR17, TCP23, HB34, and GLYMA.06G314400 were considered for *A. thaliana, G. max*, and *Z. mays.* Only these four TFs were common between species for which experimental data were available. The 3D structures of TCP23, HB34, and MYBR17 were compared for *A. thaliana* and *Z. mays.* The results highlighted higher RMSD values of 1.238 Å, 1.29 Å, and 0.847 Å for TCP23, HB34, and MYBR17, respectively **(Table 2 and Figure 4B)**. For TF GLYMA.06G314400, the structures were compared for species *A. thaliana* and *G. max.* Here also, higher RMSD values of 1.18 Å was recorded. These result indicated divergence between species for these TFs. The experimentally validated as well as PTF-Vāc generated binding motifs of these four TFs were further compared. The comparison of TFBSs between species for these TFs showed difference for both experimental as well as PTF-Vāc generated TFBSs. As shown in **Table 2 and Figure 4B**, the motif comparisons of TFs between species using TOMTOM found no significant similarities (p-value >>0.05) for both experimental and PTF-Vāc generated motifs, though there were significant agreements between experimetnally reported and PTF-Vāc generated motifs within species. This result showed that the binding motifs of TFs vary across different plant species and the developed approach was capable to learn this shift. All these series of validation and benchmarking studies reinforced that PTF-Vāc performed consistently in superlative manner and may be very useful for cross-species TFBS identification without any dependency on predefined models of TFBS, nor its performance is related to the volume of data. PTF-Vāc’s substantial advantage is also due to its pioneering use of generative and interpretable AI to learn from the co-variability of transcription factors and their binding sites, a capability that sets it apart from other methods. It can be employed successfully to even define the binding motifs in newly sequenced plant genomes.

**Table 2:**
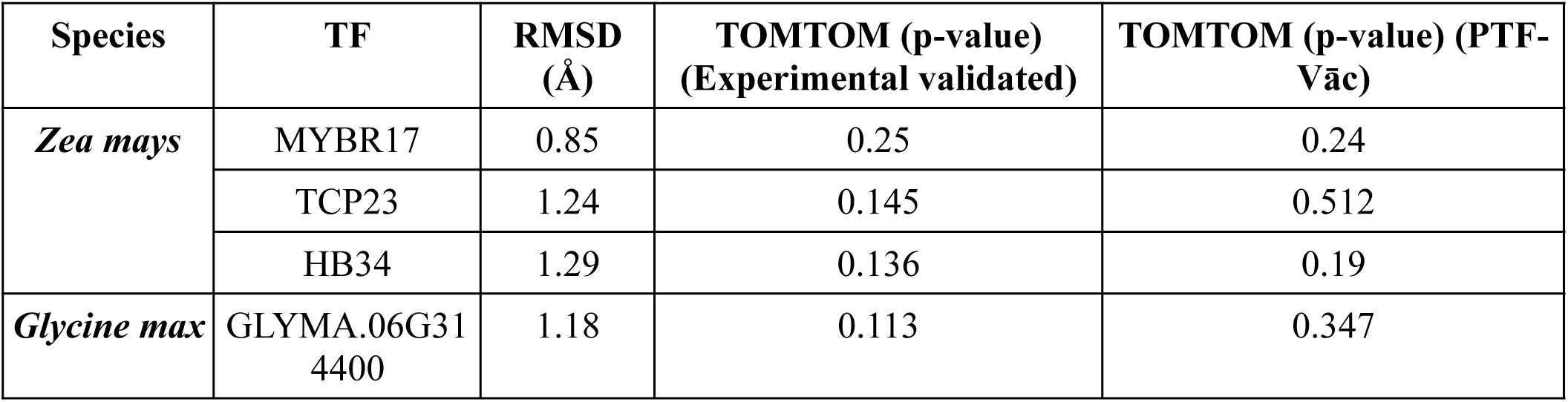
RMSD values and TOMTOM p-values for TF 3D structures and binding motif, respectively, across different plant species *A. thaliana*, *G. max*, and *Z. mays*.

In a recent benchmarking study incorporating large number of TFBS discovery tools, PTF-Vāc was found outstanding across all parameters (Jyoti et al., 2024).

### The curious case of splice variants of transcription factors and their differential binding preferences

This part of the study brings into the notice a very interesting condition of finding the binding sites of splice variants of TFs, a question vastly unaddressed in plants. ARF8 has two splice variants in *Arabidopsis thaliana* but the binding motif is defined only for one variant. For this TF, none of the existing software tool has a binding model, nor any sufficient binding experimental data which is available to detect motifs using the traditional approaches. This holds true for both of its splice variants. One completely misses out the binding site detection for the second splice variant of ARF8 as for it there is no binding data in sufficiency to raise its model. Since the existing tools rely upon the traditional motif detection approaches, they too are unable to identify the splice variants specific binding even within the same species. This means that with the existing software pool one is far away from properly understanding how various splice variants of any TF work and create spatio-temporal regulatory system for plant’s growth and development. The case of ARF8 transcription factor discussed here very aptly underlines the same.

To evaluate the efficacy of PTF-Vāc to help in such situation, we retrieved the experimental data from Ghelli et al. in 2018 (Ghelli et al., 2018). In their study, two splice variants of ARF8 exhibited distinct tissue-specific functional roles. ChIP-qPCR results demonstrated that in one scenario, ARF8.2 regulated the IAA19 gene, thereby influencing the auxin pathway. While in a different condition (later flower developmental stages), ARF8.4 regulated both the IAA19 and MYB26 genes, controlling tissue elongation and lignification. In order to validate PTF-Vāc’s capability in detecting splice variants’ differential binding sites, we initiated a protein structure similarity analysis by superimposing ARF8.2 and ARF8.4, revealing a significant structural difference with RMSD of 1.2 Å. Subsequently, PTF-Vāc was employed for these two variants (**Figure 5**). PTF-Vāc exactly identified the same two genes as the targets of these two variants of ARF8 which were reported experimentally also. The analysis indicated that ARF8.2 specifically bound to the promoter region of the IAA19 gene. However, ARF8.4 exhibited binding to both IAA19 and MYB26 genes, and the bindings sites of these variants also differed, substantiating founding assumption of PTF-Vāc algorithm that structural changes in transcription factors lead to alterations in its DNA binding sites. This study also highlights the capability of PTF-Vāc to distinguish between splice variants differential binding for any given TF which most of the existing tools are unable to answer.

**Figure 5:**
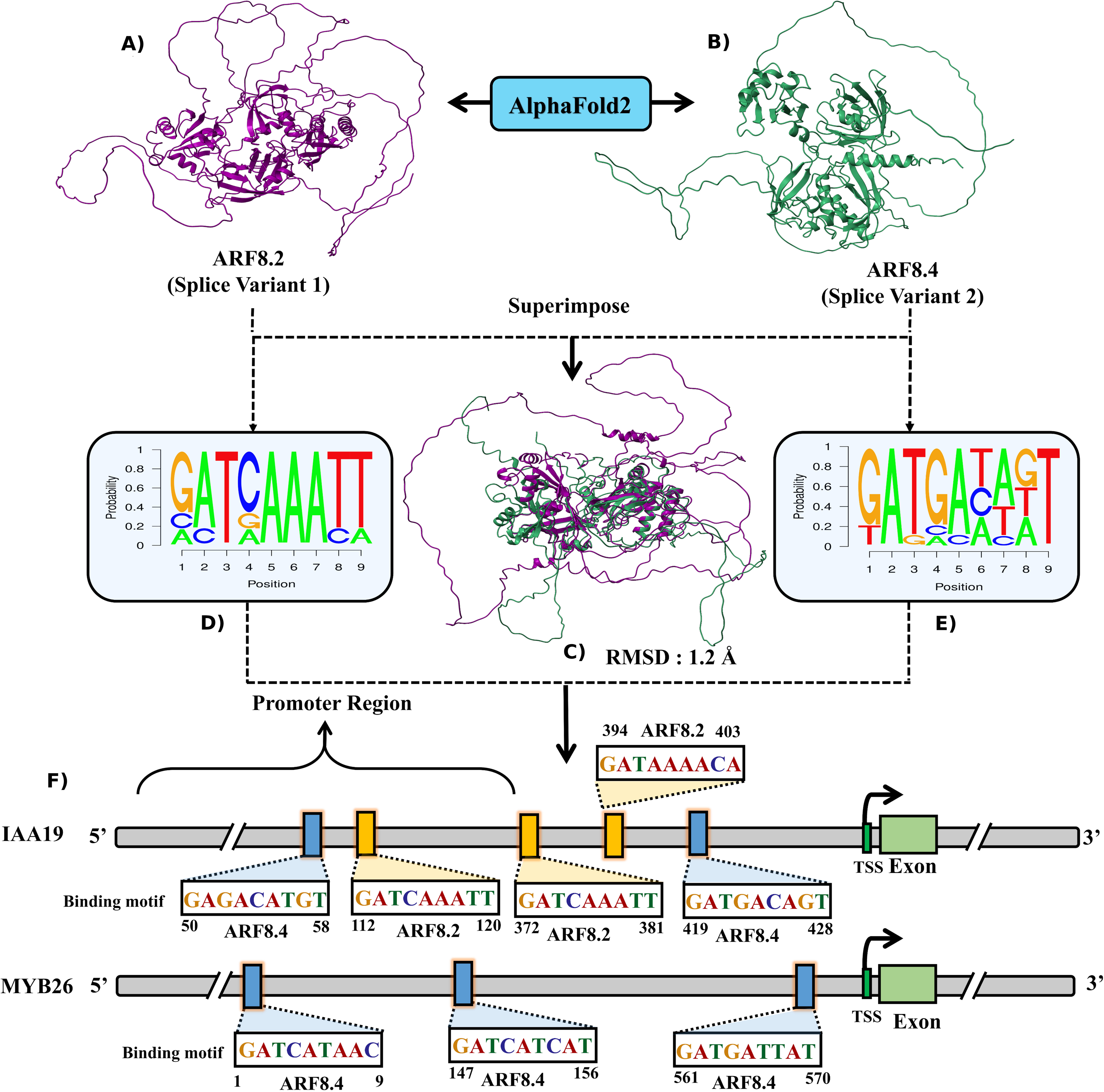
Structural Superimposition and binding sites of ARF8 Splice Variants. **A-B)** Present the 3D structures of ARF8.1 and ARF8.2 splice variants, respectively. **C)** The structural superimposition underscores significant differences between these variants with 1.2 Å RMSD. **D-E**) Motif logos for ARF8.2 and ARF8.4 illustrate distinct binding patterns. **F)** Binding sequences reveal the interaction of each splice variant with transcription factor binding sites on the promoter regions of IAA19 and MYB26 genes. Color-coded sequences emphasize variant-specific binding patterns. This figure provides a clear insight into the structural disparities and differential binding sites interactions of splice variants of a TF even within the same species, which most of the existing software fail to detect.

### Genome-scale analysis of TF binding sites in the Tea Genome using PTF-Vāc

To further demonstrate PTF-Vāc’s genome wide application, we used it to identify TF-binding sites across the entire tea genome (*Camelia sinensis*), selecting BES1 as the TF of interest. BES1 was chosen because it has been characterized through ChIP-seq in tea (NCBI project ID: PRJNA1206779), providing a dataset for validation, while the full scope of its target genes in this species remains unexplored. PTF-Vāc was expected to resolve that. PTF-Vāc was focused across the 2kb upstream promoter regions of all the genes in the tea genome, as these regions are critical for TF-mediated gene regulation. Using the 3D structure of BES1, generated by AlphaFold2, PTF-Vāc was run. It identified 1,878 genes as the potential targets of BES1.

*Validation with ChIP-seq data*.

To assess the accuracy of PTF-Vāc’s, we obtained and preprocessed ChIP-seq data for BES1. Analysis of the ChIP-seq peaks revealed that 539 genes were directly targeted by BES1. Comparing these with PTF-Vāc’s identifications, we found a remarkable overlap: 519 of the 539 ChIP-seq identified target genes (98%) were also identified by PTF-Vāc.

This high concordance underscores the reliability of PTF-Vāc in identifying biologically relevant TF-binding sites on a genome-wide scale, that too for a completely unknown species and its TF.

#### Motif discovery and comparison

We performed motif analysis on both the PTF-Vāc identified and ChIP-seq derived binding sites. The enriched motifs were then compared using TOMTOM, yielding a very significant score (<<0.01), which indicates high similarity and statistical significance between the motifs identified by both methods **(Supplemental Figure 1)**. A very interesting point to note is that the motif reported by the authors in that wor k (Liu et al., 2025) on their ChIP-seq binding data, was reported by MEME. The motif reported by PTF-Vāc on genome-wide run matched significantly to the motif reported by the authors on their ChIP-seq binding data. But the free energy of binding of PTF-Vāc motif for binding stability was found much higher than the motif reported by the authors for ChIP-seq data using MEME. Thus, this analysis presents and reiterates that how credibly genome wide runs for any TF and any species can be done using PTF-Vāc.

#### Biological relevance of BES1 Target genes

To investigate the biological significance of the 1,878 BES1-targeted genes identified by PTF-Vāc, we conducted functional enrichment analysis using gene ontology (GO) and KEGG pathway databases. The results, visualized in the added **Supplemental Figure 2 and Supplemental Table 3**, revealed significant enrichment in stress response pathways. Specifically, GO analysis highlighted terms such as “response to drought stress” (FDR = 1e^-22^) and “response to salt stress” (FDR = 1e^-16^). KEGG analysis further identified pathways related to abiotic stress signaling, such as the MAPK signaling pathway, consistent with the GO findings.

These results align with the known roles of BES1 homologs in *Arabidopsis*, such as BZR1 and BEH4, which regulate stress responses. For instance, BZR1 has been shown to enhance drought tolerance by modulating stomatal closure and stress-responsive gene expression (Cui et al., 2019), while BEH4 contributes to salt stress tolerance by regulating ion homeostasis (Feng et al., 2023). Our findings suggest that BES1 in tea plays a role in mediating abiotic stress responses, offering novel insights into its regulatory functions in this economically important crop which could be made possible due to PTF-Vāc genome wide run. This entire tea genome-scale analysis of BES1 is hosted on our web-server platform (https://scbb.ihbt.res.in/PTF-Vac/Application/).

### Web-server implementation

The PTF-Vāc server hosts the software in a user friendly manner for the identification of TFBSs within plant DNA sequences. To identify TFBS, users are required to upload a genomic DNA sequences in FASTA format and its associated TF in “pdb” format 3D structure generated from Alphafold2. The experimental 3D structures for most plant TFs are unavailable while PTF-Vāc needs just relative structural information where precise structural information is not necessary but uniform source is important. It is important to note that PTF-Vāc, unlike the existing tools, is capable of discovering binding sites irrespective of the number of input sequences, performing well even with a single input sequence. It remains immune to the noise in input data. Upon uploading the “pdb” file, the user can instantly visualize it on the home page. Once the user submits the data, the backend processing begins with the PTFSpot which plays a critical role by initially identifying potential TF binding regions within the uploaded DNA sequences specific to the given TF. These identified binding regions are then passed to the core functionality of PTF-Vāc to generate actual TFBS motifs. The results are presented to the user in a comprehensive and interactive format on a dedicated results page. Users also get the capability to download the results in a tabular format, facilitating further downstream analysis and integration with other tools. PTF-Vāc’s output for a given input sequence is the most informative site, the most potential binding sequence/motif. For scenarios involving multiple DNA sequences, PTF-Vāc offers features like as: motif logo which is a graphical representation of the identified TFBS, position weight matrices, and consensus representation which can be used and compared. The results page also provides explainability scoring plot in form of Grad-CAM score while highlighting the generated motif regions in the sequence.

Grad-CAM helps users understand which sequence context features have the strongest impact on binding status for each motif. It identifies significant k-mers crucial for binding activities and the top-scoring k-mers often closely match the reported core motif of that TF. The Grad-CAM interpretability serves as a strong confidence measure for the proposed TFBS motifs. This provides users with additional independent confidence and insights into other potentially important regions found critical for TF binding. Another important feature of PTF-Vāc is its integration with TOMTOM in the backend. This allows users to compare the PTF-Vāc generated motif’s PWM with their own choice of PWM, enabling a comparative analysis of binding motifs across different datasets or experimental conditions. Such functionality enhances the flexibility and utility of PTF-Vāc in diverse research contexts.

To enhance the utility of the web-server for genome scale studies, it has been integrated with pre-loaded genomes in GFF3 format of annotations along with annotated promoter regions of their genes (e.g., ±2 kb from TSS) into the web server. At the present the server has genomes for four model species viz. *Arabidopsis thaliana*, *Oryza sativa*, *Glycine max*, and *Zea mays*, for which completed full genome along with corresponding Alphafold2-derived PDB structures are available. In the future, more genomes may be integrated, depending upon their completion, and availability/generation of corresponding PDB structures, a continuous process in overall.

In addition to all, the server result page also provides additional filters and scoring schemes for the users. The implemented scoring schemes are as following: **i) Scoring Scheme Implementation:** a binding site confidence score based on the Grad-CAM based importance score profiles to identify the sequence context features having strongest impact on binding status for each motif. This score reflects the likelihood of each generated binding site being a important binding site, allowing users to rank these binding sites accordingly (**Figure 6**).

**Figure 6:**
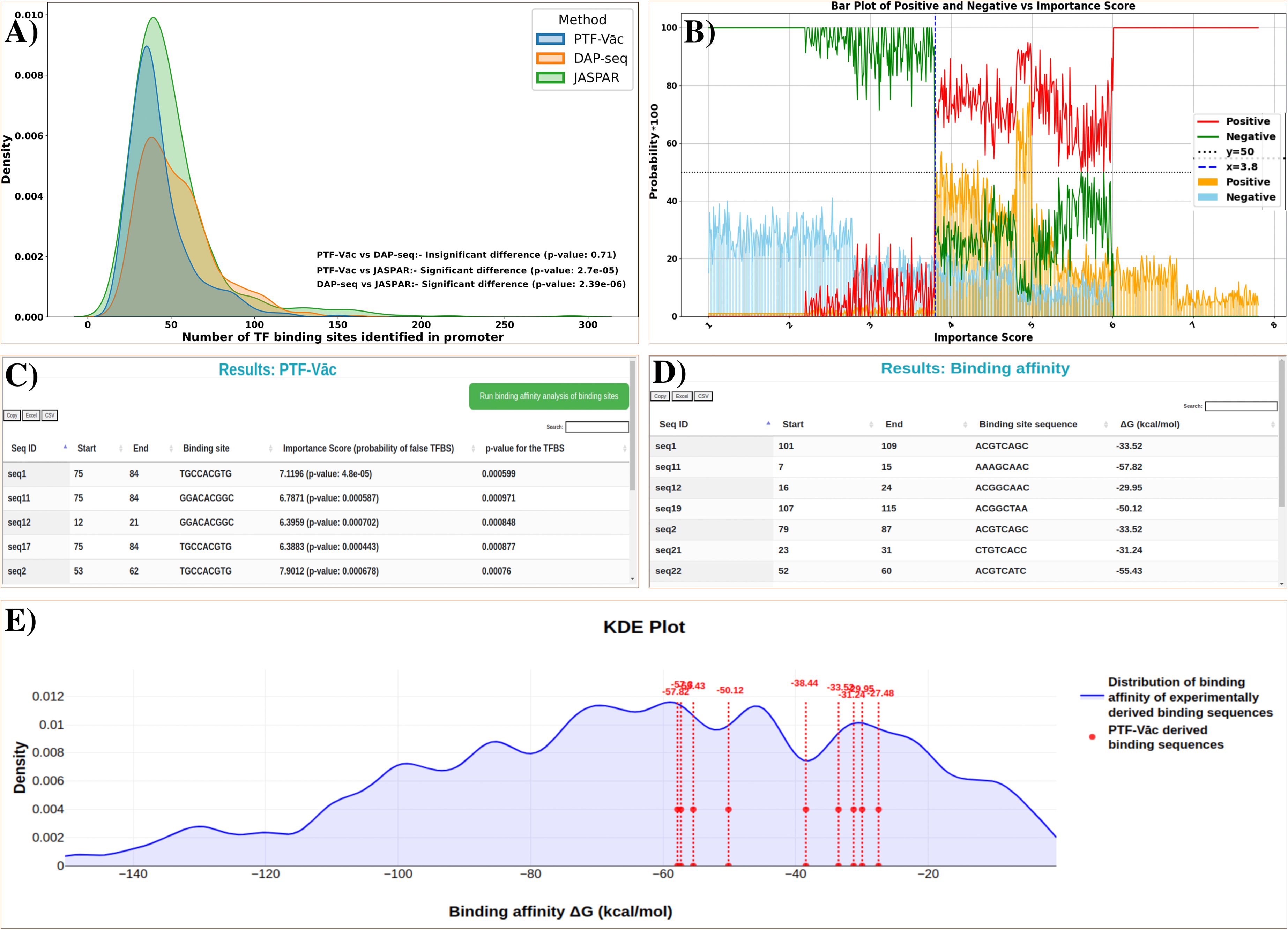
**A)** The number of TF binding sites identified by PTF-Vāc, DAP-seq, and FIMO using JASPAR data in promoters of *Arabidopsis thaliana*. **B)** The probability of binding sites being positive or negative across importance score intervals. The plot quantifies the probability (Y-axis) that a TFBS site functions as a positive or negative binding site event within binned importance score ranges (X-axis). PTF-Vāc server uses this distribution to assign the p-value confidence scoring for the importance scores, in its result page for every proposed TFBS. **C)** Snapshot providing description of new improved result table with importance score, its binomial test based p-value, and hypergeometric test based p-value for every binding sites identified. **D)** Snapshot providing description of molecular docking analysis using HADDOCK3 output in tabular form which includes binding affinity between TF-3D structure and its binding sites. **E)** Snapshot depicting KDEplot of binding affinity values from molecular docking analysis for the set of 563 TF-3D structure and their binding sites.

To determine a threshold, a density plot of Grad-CAM based importance scores has been used in the back-end, having positive and negative binding sites distributions plotted against scores ranging from below 1 to 8 **(Figure 6B)**. The analysis revealed distinct patterns i.e., **i) Scores below 3.8:** These are predominantly associated with negative binding sites, exhibiting a high probability of being negative (ranging from 0.7 to 1.0), while the probability of being a positive binding site is low (0 to 0.25). **ii) Scores above 3.8:** experimentally validated positive binding sites are enriched within this range, indicating a higher likelihood of biological relevance. **Figure 6C** illustrates how these scores and their probability of being a false TFBS is listed in the result page. Within the score range of 3.8 to 8, the probability of a site being a positive binding site is >0.7. To ensure the reliability of this threshold, a t-test to compare the distributions of Grad-CAM scores for positive versus negative binding sites yielded a significant p-value (p < 0.05), confirming that the score distributions differ meaningfully between these two groups and supporting the use of 3.8 as the suitable cut-off.

For users requiring greater stringency, the threshold can be adjusted upward. A binomial test based p-value scoring is also implemented for further support **(Figure 6C)** the observed importance score.

**i) ii) Enrichment test:** A hypergeometric test was implemented for all the identified binding sites with their p-values, given in the output/result page of the web-server (**Figure 6C**) which suggests the chance of seeing the observed TFBS in any random data.
**ii) Binding affinity analysis:** The web-server has also implemented HADDOCK3 (Giulini et al., 2025) for TF-Binding site molecular docking analysis, where a user can find the binding affinity for the proposed binding site with the corresponding TF (**Figure 6D**). In addition, we have also done a comprehensive molecular docking analysis comprising of 563 experimentally validated position weight matrix (PWM) of TF binding motifs and their corresponding TFs. The 3D structures of DNA motifs corresponding to each transcription factor were generated using the DNA sequence to structure web server (https://scfbio-iitd.res.in/software/drugdesign/bdna.jsp). Corresponding Alphafold2 generated TF 3D structures for every binding motifs were used. Protein-DNA MDs were then performed to get the binding affinity scores between TFs and their corresponding binding DNA motifs using HADDOCK3 (Giulini et al., 2025). The docking results based scoring distribution is presented as a KDEplot on the same page where the binding affinity scores are displayed (**Figure 6E**). So, the user can also visualize if the detected binding sites are thermodynamically supported and falls in the range of experimentally proven sites binding free energy or not.

PTF-Vāc is built using PHP, JavaScript, and HTML5 for frontend development, ensuring a responsive and intuitive user interface. The backend processing leverages from the Apache-Linux platform, with significant components developed in Shell and Python scripts. The project was executed within the Ubuntu Linux platform’s Open Source OS environment. **Figure 6 and 7** provides an overview of PTF-Vāc web-server implementation. The PTF-Vāc system is freely accessible at https://scbb.ihbt.res.in/PTF-Vac/. The entire software is also freely available (available at: https://gitlab.com/scbblab/ptfvac) as standalone version along with source code, using which one can run it locally on huge volume of inputs and large genomic sequences.

**Figure 7:**
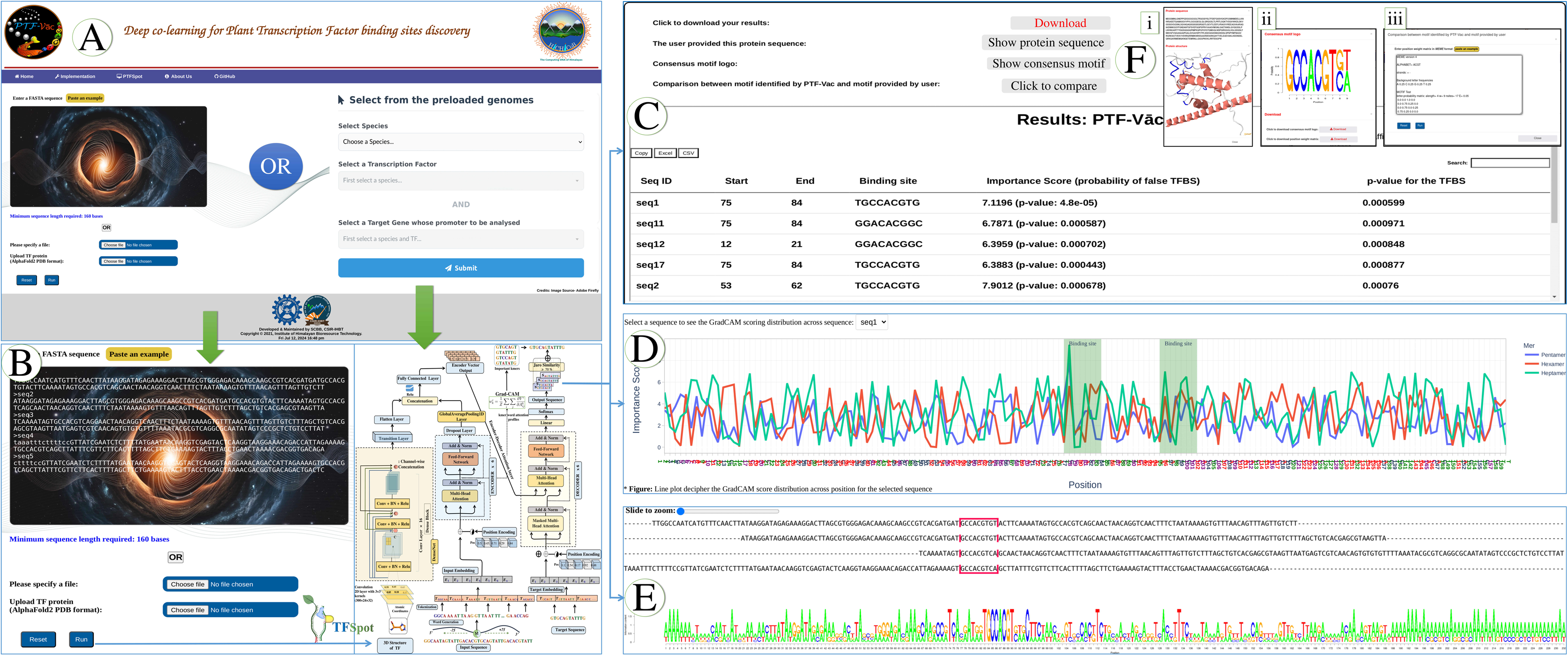
PTF-Vāc webserver implementation. **A)** Input data box where the user can either paste or load the input files or the user can directly select from the pre-loaded genomes with annotated promoter regions (e.g., ±2 kb from TSS) with corresponding Alphafold2-derived PDB structures from the dropdown menu, **B)** Inputs are genomic DNA sequence in FASTA format and TF 3D structure pdb format generated from Alphafold2. **C)** The binding site information is represented in the form of table. Download option for the result in the tabular format. **D)** Also, important feature scoring is implemented and is represented in the form of interactive line plot depicting the distribution of scoring distribution across the positions for the selected sequence. **E)** The binding site is highlighted among the sequences and a motif logo is also built from consensus. **F)** User can visualize 3D structure of protein with its sequence, sequence logo of the binding sites and can download its PWM.

### Limitations of PTF-Vāc

**i) Requirement for 3D TF Protein Structures:** While the tool leverages from AlphaFold2 3D structures because experimental 3D structures for most plant TFs are unavailable, the availability of a reliable 3D structure for the specific TF is a mandatory input for PTF-Vāc to perform at its best. Generating 3D structure from Alphafold2 may consume some additional time.
**ii) Limited to Plant TFs only**: While PTF-Vāc is a universal model for TF:DNA interactions, its development, training, and extensive benchmarking studies are exclusively focused on plant TFs and plant genomes. The challenges it specifically addresses, such as high cross-species variability and context-dependence of TF:DNA interactions in plants, are particular to this biological domain. Therefore, the proven utility and performance of PTF-Vāc are restricted to the plant kingdom. This strategy of PTF-Vāc can be extended further to animal kingdom.
**iii) Condition-Specific TF Binding:** Condition-specific TF binding refers to the phenomenon where TFs bind to DNA in a cell type-, tissue-, developmental stage-, or stimulus-dependent manner. This dynamic binding allows cells to precisely regulate gene expression in response to internal and external signals. PTF-Vāc can tell the most potent binding sites for any given TF, but it does not tell about the condition specific binding. Condition specific binding is a highly multi-factor dynamic network based phenomenon, posing a completely different set of problems to be solved.

In summary, cross-species variability is a pronounced feature of plant genomes which is duly reflected across its various components including TFs and their binding sites. This means that the age old practice of using TFBS models and matrices developed for model species for TFBS finding in other species is flawed and a big source of misinformation. Most of the existing tools are coupled to the process of defining a TF specific binding matrix/motif/model using experimental binding data for the given TF for the selected species, as their first step. This way they do not address cross-species variability and are not even able to answer the differential binding preferences of the splice variants of any TF even within the same species. The present work has put forward an innovative approach, PTF-Vāc, which has learned the co-variabilities of TF structure and its associated binding sites preferences while utilizing the universal TF:DNA interaction model of PTFSpot. PTF-Vāc uses deep-learning encoder-decoder generative systems to speak out the binding sites in any given DNA sequence in completely *ab-initio* manner. Its output is also supported by DL-Interpretability scheme. This all makes it completely free of the shortcomings of the existing software pool and from any dependence on predefined TF specific models and motifs. Thus, it is capable to accurately identify the binding sites of even totally novel TFs and across newly sequenced genomes even in total lack of TF information and any prior information. Thus, it is going to have a profound impact in the area of plant science and regulatory research as with PTF-Vāc, one would not need to carry out binding experiments to generate the binding data and identify the binding motifs, and then run them for whole genome scanning. Nor one would be anymore compelled to use TFBS models from other species to find the binding preferences in some other species, an age old wrong practice. And even the differential binding preferences of splice variants of transcription factors within the same species can now be detected accurately. It will need just a simple run at PTF-Vāc server with the user provided DNA sequences and the Alphafold2 derived protein 3D structure whose binding site the user would like to know across some genomic sequence.

## Materials and Methods

### Dataset retrieval

For the development of universal and generalized model for the identification of TF binding sites, we retrieved data for *Arabidopsis thaliana* from our previous work (Gupta et al., 2024), where we created six different datasets using ChIP/DAP-seq. The positive instances of binding regions for any TF and its corresponding 3D protein structure associated with each TF were extracted from Dataset “E” while maintaining the methodology introduced by PTFSpot (Gupta et al., 2024). A total of 40 TFs representing 40 different families were considered for model training. By considering this subset of TFs, we could effectively represent the entire set of 325 TFs in Dataset “E” without imposing excessive computational demands. In this dataset, each TF represented a unique TF family, and we selected the TFs with the highest binding data within their respective families. This focused selection aimed to overcome the challenges associated with data handling and computational resource requirements. The TF structures were derived using AlphaFold2 (Read et al., 2023) as experimental 3D structure of most of the TFs of plant species are unavailable and would be so for long time. Also, for relating structural variability, all TFs’ structures derived from a common approach solves the purpose of learning the relative changes. The dataset was split in a 70:30 ratio for training and testing purposes.

### Transformer-DenseNet system

The 3D structure of a protein plays a crucial role in determining its DNA binding sites like lock and key. In our approach, the structural information of TFs for each of the corresponding binding region extracted from DAP-seq data were integrated. A DenseNet model based on 3D protein structure was raised with a Transformer encoder which takes sequences as input. This integration was achieved through a bi-modal architecture, creating a universal model capable of identifying TF binding sites in a global and cross-species manner. Previously, we had developed a 3D protein structure-based DenseNet, which utilized the spatial coordinates of all the atoms in the protein structure as input (Gupta et al., 2024). In the cases where the length of a protein was shorter, padding with zero values was done for the empty columns. A window size of 300 covered the amino acid positions, with each position having a vector with elements corresponding to the 24 atoms, each of which is mapped to three-dimensional space using normalized X, Y, Z coordinates. Each vector position signifies a specific atom number. This DenseNet system was implemented in pytorch.

### Constructing the DenseNet architecture

The DenseNet model processes a 3D tensor representing protein structure 300×24×3 (300 amino acids, 24 atoms per amino acid, three X, Y, Z space coordinates for each atom). This architecture, consisted a single convolution layer with 32 convolution filters (kernel size = 3), batch normalization, and 2D max-pooling (stride = 2), and incorporates four dense blocks and three transition layers, totaling 121 layers. Dense blocks, fostering feature reuse through dense connectivity, play a crucial role. In a dense block, each layer ***“l”*** receives direct input from all preceding layers, producing output through operations like convolution and normalization. Transition layers effectively reduce the dimensions before the next dense block.

Expressed mathematically, let ***“H_l_”*** represent the output feature maps of layer ***“I”***. The output of layer ***“I”*** is computed as:

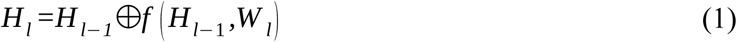

Here, “⨁” signifies concatenation, “***W_l_***” denotes the layer’s weights, and “***f***” incorporates batch normalization (BN), rectified linear units (ReLU), and convolution operations. The unique dense connectivity enhances feature reuse, contributing to the model’s ability to learn complex hierarchical features effectively.

The growth rate, denoted by the hyperparameter “***k***” in DenseNet, emerges as a pivotal determinant in the architecture’s extraordinary performance. DenseNet’s innovative design treats feature maps as a global network state, enabling impressive results even with a smaller growth rate. At each layer, “***k***” feature maps contribute to this global state, where the total number of input feature maps “***F_m_***” at the “***l^th^***” layer is calculated as:

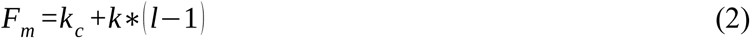

The term “***k_c_***” denotes the total number of channels in the input layer. To enhance computational efficiency, a 1×1 convolution layer precedes each 3×3 convolution layer. This design, known as a bottleneck layer, reduces the number of input feature maps, typically exceeding the output feature maps “***k***”, by generating 4k feature maps. This architecture’s output is intertwined through a bi-modal architecture in a feed-forward layer within the transformers encoder to capture TFs’ relationship with input data to raise a universal model to identify TF:DNA interactions in global and cross-species manner. For a more in-depth understanding of the implementation of DenseNet architecture, additional details can be found in our previous study (Gupta et al., 2024).

### Word representations of sequence data for Transformer

Each sequence can be considered as a collection of independent words. Every occurrence of the dimers, trimers, tetramers, pentamer, hexamer, and heptamer sequences was transformed for the verbal representation of the sequences in an overlapping window. All these kmers provide information on base stacking, shape, and regional motif information (Zhou et al., 2013; Sharma et al., 2021; Hombach et al., 2016). Transformers takes two input i.e., source and target input. The source input vector had a maximum length of 465 words and a sequence length of 159-162 bases.

The target input vector which was motif for that particular sequence had a maximum length of 12 words and a sequence length of 9-12 bases. Also, target sequences contains two flanking words situated adjacent to the binding sites. Distinct integers were used as a token for each unique word. The tokenized version of the sequences were converted further into numeric vectors and matrices, known as embedding. The transformer encoder and decoder fed the tokenized and embedded sequences as input to train and assess the models. The TensorFlow (Keras) tokenizer class was used to implement the tokenization procedure.

### Implementation of the Transformers Encoders-Decoders

The encoder-decoder architecture in Transformers is crucial for handling sequential data. The encoder processes the input sequence, extracting hierarchical features, while the decoder generates the output sequence by attending to relevant parts of the input sequence. This bidirectional flow of information ensures the model’s ability to capture dependencies and relationships within the input and generate contextually rich sequences as output. The integration of attention mechanisms in both the encoder and decoder enhances the model’s capacity to handle transcription factor motif translation.

### The Encoder

Encoder implements a multi-headed attention mechanism which in itself comprises self-attention layers, which allow the encoder to attend to different words of the input and consider their relationships when making decisions. The self-attention layers use dot-product, allowing the model to weigh the importance of different input elements based on their relationships. This enables the transformer to capture the input data’s long-range dependencies and contextual relationships. The sequences from the previous step’s tokenized representation were fed into the input layer of the Transformer system. Input sequences were padded to ensure a constant size of 465 words. To create an output matrix formatted as samples, sequence length, and embedding size, embedding converted each word’s token into a word vector whose length equals the embedded size.

The input sequence of word tokens is denoted as *“**Z**”*, with a length of *“**I**”*. The embedding layer projects these discrete input tokens into continuous vector embeddings. If “***v****”* is the size of the vocabulary and “***d****”* is the dimensionality of the word embeddings, the embedding layer is represented as a matrix 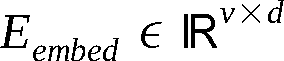.

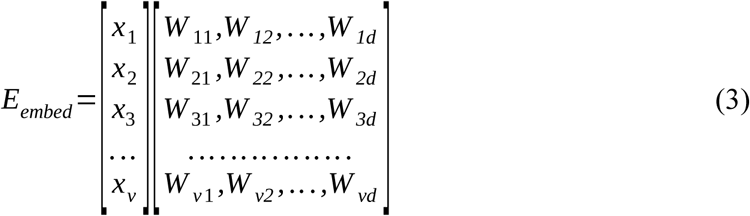

Each word in matrix “***E_embed_***” combines with its positional embedding “***P****”*, where “***P****”* shares dimension “***d****”* with the word embedding vector. The resulting matrix, “***E_embed_***”, is obtained through “ *E’_embed_ =E_embed_ +P*”, where “***P****”* is calculated using sinusoidal positional embedding equations.

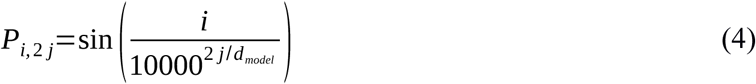

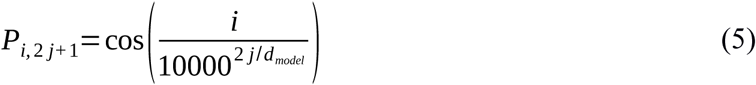

Here, “***i***” represents the position of the token, and “***d_model_***” is the embedding dimension. “ *E’_embed_*” matrix enters the transformer encoder block, which processes it through a multi-head attention layer. This module involves multiple heads, each dividing its query, key, and value parameters N-ways and independently processing the splits. The output from each head produces an attention score derived from the key, query, and value computations.

The process involved the repetition of the same steps, where individual attention vectors were generated and then concatenated. These concatenated vectors were subsequently forwarded to the feed-forward network block within the encoder for further processing. For further insights into the details of the encoder section and the mathematical equations governing the self-attention mechanism, it is advised to refer the PTFSpot work (Gupta et al., 2024). As a measure to mitigate overfitting, the output from the multi-head attention layer was supplied to a dropout layer of 0.1 fraction, succeeded by a normalization layer. Following normalization, the output was directed to fully connected feed-forward layers with 64 nodes that converges in the subsequent dropout layer of 0.1 fraction, succeeded once again by a normalization layer. After normalization, the output is seamlessly fused with the DenseNet output as discussed above, encapsulating vital structural information from the 3D protein structure. This amalgamated result is channelled into fully connected feed-forward layers, converging within the subsequent dropout layer and followed by another normalization layer. Subsequently, this enhanced output from the encoder integrates into the decoder block of the transformer.

### Decoder block of the transformer

The decoder block mirrors the structure of the encoder, comprising three sub-layers: Masked Multi-Head Attention, Multi-Head Attention, and Feed-forward Network. Notably, there are two Multi-Head Attention sub-layers in the decoder, with one of them being masked. Each decoder being supplied with two inputs: one originating from the target sequence, and the other one derived from the encoder’s representation. This intricate interplay significantly enhances the decoder’s capacity to assimilate information from both its preceding counterpart and the comprehensive encoding of the input sentence by the encoder. The Masked Multi-Head Attention mechanism in the decoder plays a pivotal role in capturing dependencies within the target sequence while preventing information leakage from future positions during training. This attention mechanism allows the decoder to attend to different parts of the input sequence, aiding in the generation of contextually rich representations for each position in the target sequence.

The Transformer architecture implements Masked Multi-Head Attention in the decoder through a series of mathematical operations. Consider the target input sequence as “***X_T_***” and its embedding as “***E****”*, where “***E****”* is the output embedding matrix obtained after converting the input into word embedding and adding positional encoding, similar to encoder block before being fed into the Masked Multi-Head Attention layer.

### Masked Multi-Head Attention of Decoders

For each distinctive head “***i***”, symbolizing “***i=1, 2,…., h,”*** the calculation of three pivotal matrices ensues: ***Qi*** (Query), ***K_i_*** (Key), and ***V_i_*** (Value). This is accomplished through the matrix multiplication of “***E***” with the corresponding weight matrices “***W_qi_***”, “***W_ki_***”, and “***W_vi_***”. The formulation of attention scores “***Z_i_***” transpires through the application of the softmax function to the scaled dot-product attention mechanism, adhering to the expression:

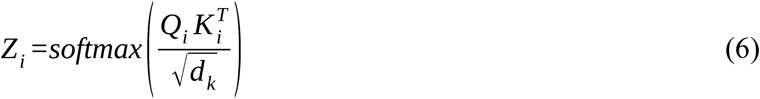

To preserve causality during training, a masking mechanism is applied to the attention scores, specifically designed to impede the model from attending to future positions, encapsulating the essence of masked self-attention. The ensuing phase involves the computation of the weighted sum, where each head “***i***” contributes to the formation of “***Head_i_***” via the formula:

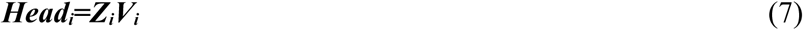

A concatenation of the outputs from all heads transpires, culminating in a subsequent multiplication with the output weight matrix “***W_o_***”, yielding the definitive output of the Masked Multi-Head Attention mechanism in the decoder:

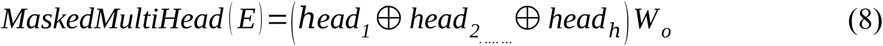

This exposition encapsulates the procedures involved in applying masked self-attention across multiple heads within the decoder framework. In the decoding phase of a Transformer model, after the Masked Multi-Head Attention mechanism processes the input sequence, the encoder output is incorporated as the second input for the decoder. The Decoder Multi-Head Attention mechanism is then employed, similar to the encoder, enabling the model to selectively focus on relevant parts of the input sequence and the encoded information. Subsequently, the output from the Decoder Multi-Head Attention mechanism undergoes further refinement through the Feed-forward layer. In decoder processing, layer normalization and residual connection enhance information flow, stabilizing the network. Subsequently, the final output undergoes conditional probability estimation, unfolding as a crucial aspect of the decoding mechanism. Building upon the output after layer normalization, the model proceeds to compute the conditional probability distribution over the vocabulary for the subsequent token (Wu et al., 2022). This involves employing the softmax function, transforming raw scores into a probability distribution. The resultant distribution guides the selection of the next token in the sequence, a pivotal step in generating contextually meaningful and coherent outputs. The conditional probability *P* ⟨ *y_t_*|*y*_1_,….*,y_t_* _−_*_1_, x* ⟩ for the next token “***y_t_***” is computed using the following equation:

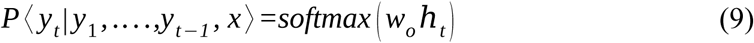

Where, “***w_o_***” is the weight matrix, and “***h_t_***” is the hidden state at position “***t***”. This process repeats iteratively for each time step until the entire sequence is decoded. The model uses the generated tokens as input for subsequent time steps, allowing for autoregressive decoding. In the context of evaluating the Transformer model’s decoder performance, the BLEU score serves as a crucial metric (Papineni et al., 2002). The BLEU score measures the similarity between the model-generated sequences and target sequences, providing a quantitative assessment of the quality of the generated output.

The BLEU score is calculated based on the precision of 5-grams in the generated sequence compared to the reference sequence. The formula for BLEU score is given by:

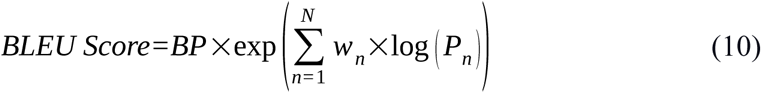

where, “***BP***” is the Brevity Penalty, accounting for the length of the generated sequence relative to the reference. “***N***” is the maximum order of 5-grams considered. “***w_n_***” is the weight assigned to each n-gram precision. “***P_n_***” is the precision of 5-grams in the predicted sequence. This formula quantifies the precision of the model’s output against reference sequences, considering 5-gram orders and assigning appropriate weights.

### Generating binding sites from the Transformer-Decoder output

The decoder segment of the transformer is designed to generate multiple words (5-mers, 6-mers, 7-mers) as part of its output sequence comprising both the motif and flanking bases for each provided sequence. These multiple words are supposed to map to the target (the binding motif) in an overlapping manner where each such word aggregating to a common region in overlaps add to the confidence. The same is repeated for the 5’ and 3’ flanking regions for a single 7-mer words to act as the anchors for the binding region and boost the collective confidence value for the detected target region where three such different regions come in support, collectively for a single region and with multiple spoken words. The crucial step post-decoding involves matching these generated words with target sequences. The goal is to identify the most suitable sequence region based on a similarity threshold. For this task, the Jaro similarity algorithm (Basak et al., 2023) was applied. Jaro similarity is a string matching algorithm that give a similarity value between 0 and 1. A value of 0 indicates no match, while 1 signifies a perfect match. The words generated by the decoder are individually matched with the target sequence for its every position in a sliding window manner, and their Jaro similarity scores are calculated. The region with the highest aggregated Jaro score with highest aggregations of the words is identified as a potential binding site component. The Jaro Similarity “ *J* _(*S S j*)_” between two strings “***S_i_***” and “***S_j_***” is computed using the following equation:

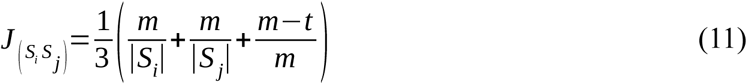

where, “***m***” is the number of matching character, “***t***” is half the number of transpositions (characters that match but are in the wrong order), “***S_i_***” and “***S_j_***” are the lengths of the two strings.

Likewise, the anchoring word or the flanking bases are mapped back to the sequence to determine its anchoring position while calculating Jaro score. In cases where multiple positions were identified within the sequences, preference was given to the position where the flanking anchor words mapped besides the identified binding site as well as the Jaro similarity was calculated max. This process enhances the model’s ability to generate meaningful and contextually aligned binding site sequences for transcription factor, showcasing the power of encoder-decoder deep-learning system for speech translation from a long DNA sequence to sequence generation of a precise binding motif. The resulting final model was saved in **“*pt*”** format. The whole Transformers-DenseNet system was implemented in pytorch except the tokenization procedure which was implemented in the TensorFlow (Keras) tokenizer class. **Figure 1** depicts the operation of the implemented deep-learning encoder-decoder system.

### Implementation of Grad-CAM

Significant efforts are underway to enhance the transparency and interpretability of deep learning. DLs perform exceptionally good but are hard to explain the causality. Interpretability of DL models has been the frontier of DL research at present. It is crucial to make deep learning models more interpretative. Selvaraju et al., 2017, introduced the Grad-CAM technique, which offers a visual explanation of deep learning models (Selvaraju et al., 2017). As the deep learning models become increasingly complex, the need for interpretability grows. We implemented Grad-CAM score profiles to identify the sequence context features having strongest impact on binding status for each motif. The gradient of any target class **“c”** and the global average of the activation feature map **“*F”*** of the last attention layer ***“l”***, were computed. Then, the class-discriminative profiles of the k-mers were obtained using a weighted combination of these activation maps, followed by a ReLU function to ensure that only the features with a positive influence are considered. The mathematical formulation and operational principles of Grad-CAM are described as follows:

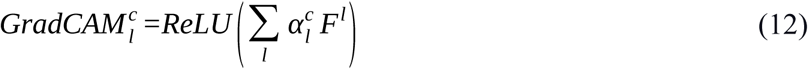

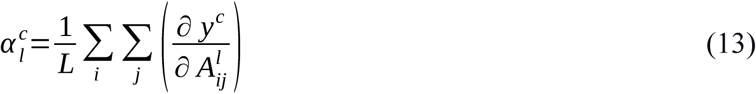

Where, 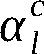 denotes the neuron importance weights, highlighting the most relevant features that contribute to the models accuracy. **‘*L’*** is the sequence length in the feature map, and 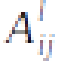 represents the activation at ***ij^th^*** position in the sequence.

In the implementation of Grad-CAM in PTF-Vāc, we chose the normalization layer after the multihead attention layer (MHA) as the layer of interest **(Figure 1)**. This layer contains distribution maps for various sequence k-mers. The key reason for selecting the normalization layer over the MHA output lies in how feature importance is captured. MHA produces attention weights, which indicate dependencies between different positions in the sequence. In contrast, the normalization layer operates after attention and feed forward transformations, refining the feature representations and ensuring they are more stable and meaningful for gradient-based interpretability.

## Software, Code and Data Availability

All the secondary data used in the present study are publicly available and their due references are already cited in the MS at appropriate places. The web-server is freely available at https://scbb.ihbt.res.in/PTF-Vac, while the freely available source-code and standalone version has been made available as at GitLab (https://gitlab.com/scbblab/ptfvac).

## Author’s contributions

S.G. carried out the major parts of this study. Jyoti carried out the benchmarking study. U.B., V.K., and Jyoti carried out the analysis in the study. S.G. and A.S. developed the web-server of PTF-Vāc. R.S. conceptualized, designed, analyzed, and supervised the entire study. S.G., Jyoti, U.B., V.K., and R.S. wrote the MS.

## Acknowledgements

S.G., V.K., and A.S. are thankful to DBT, India for financial support as project associateship. Jyoti is thankful for CSIR-UGC SRF fellowship. U.B. is thankful for DBT SRF fellowship. The authors are thankful to Ritu for her contribution for designing logo of PTF-Vāc. This MS has CSIR-IHBT MSID 5835.

