## Supplemental File for "PTF-Vāc: An explainable and generative deep co-learning encoders-decoders system for *ab-initio* discovery of plant transcription factor binding sites"

Palampur (HP), 176061, India.

^2^Academy of Scientific and Innovative Research (AcSIR),

Ghaziabad, Uttar Pradesh- 201002

Authors’ email addresses:

SG:

Jyoti:

UB:

VK:

AS:

RS:

*

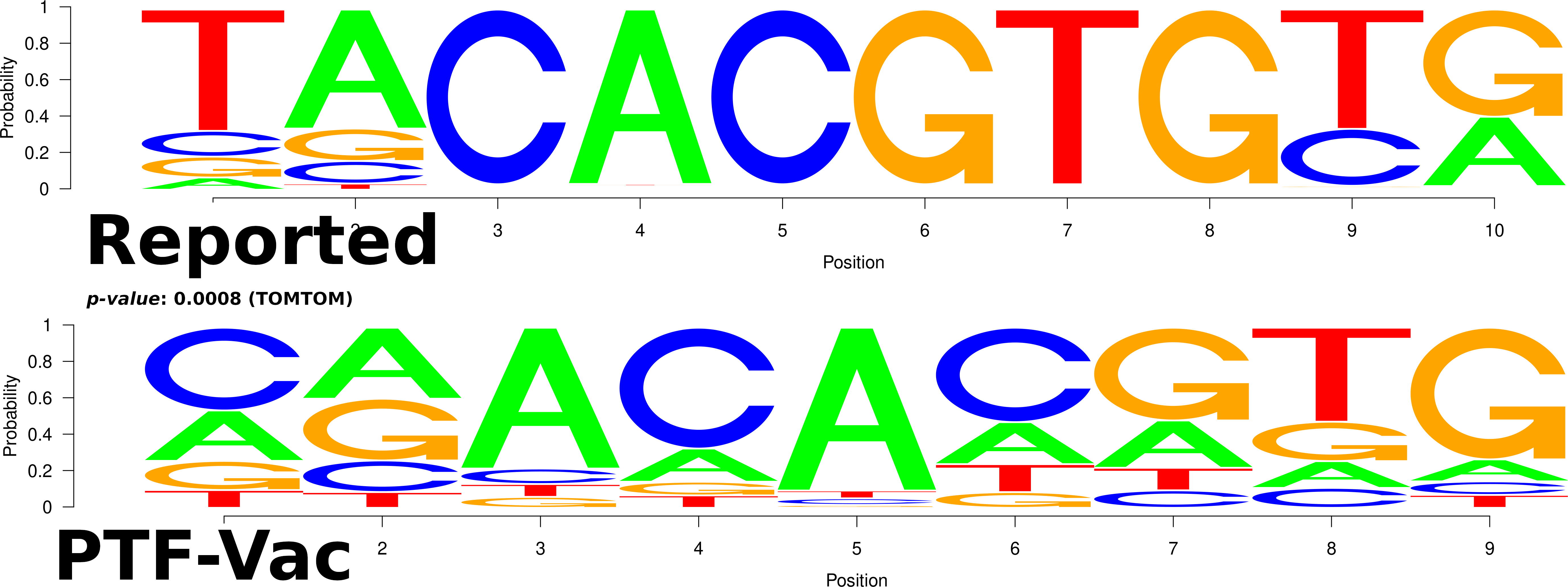
**Supplemental Figure 1:** The image displays a comparative sequence motif analysis between a reported TFBS (top panel) and a motif discovered using the PTF-Vāc model (bottom panel) for the TF BES1 in Tea. Between the logos similarity is observed between the two motifs. This similarity is further supported by a statistically significant p-value of 0.0008, obtained using the TOMTOM motif comparison tool.

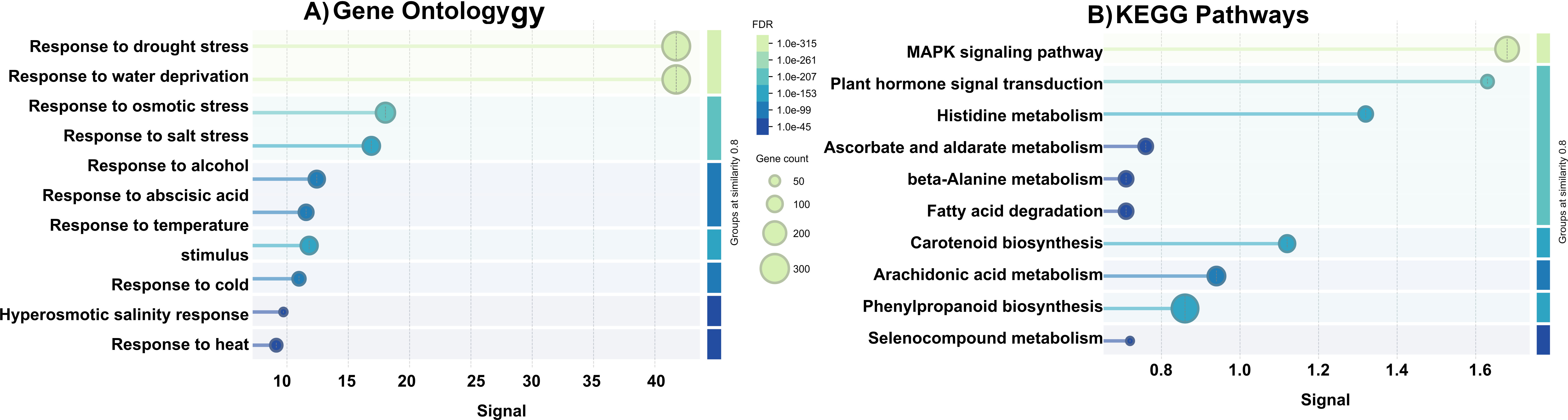

**Supplemental Figure 2:** Functional enrichment analysis of BES1-Targeted Genes. The figure illustrates the functional enrichment analysis of 1,878 BES1-targeted genes using **A)** Gene Ontology (GO) and **B)** KEGG pathway databases.

### **Supplemental Table 3: Functional enrichment analysis.**

| **Category** | **Term** | **FDR** | **Gene Count** |
| --- | --- | --- | --- |
| GO BP | Response to drought stress (GO:0009415) | 1e-22 | 55 |
| GO BP | Response to salt stress (GO:0009651) | 1e-16 | 47 |
| GO BP | Response to water deprivation (GO:0009414) | 1e-18 | 24 |
| GO BP | Response to osmotic stress (GO:0006970) | 1e-14 | 16 |
| KEGG | MAPK signaling pathway (KEGG:04010) | 1e-12 | 30 |
| KEGG | Plant hormone signal transduction (KEGG:04075) | 1e-10 | 58 |
